# CYP722A1-driven 16-hydroxylation of carlactonoic acid regulates the floral transition in Arabidopsis

**DOI:** 10.1101/2025.01.17.633504

**Authors:** Masaki Kuno, Ayumi Miyamoto, Hinako Takano, Masato Homma, Nanami Shiotani, Kiyono Uchida, Hirosato Takikawa, Masatoshi Nakajima, Masaharu Mizutani, Takatoshi Wakabayashi, Yukihiro Sugimoto

## Abstract

Strigolactones (SLs) are multifunctional plant hormones and rhizosphere signals with diverse structures, broadly categorized as canonical or noncanonical SLs. In *Arabidopsis thaliana*, SL biosynthesis mutants exhibit increased shoot branching and early flowering, underscoring their roles in developmental regulation. Shoot branching inhibition in Arabidopsis depends on the methylation of carlactonoic acid (CLA), a key intermediate classified as a noncanonical SL, catalyzed by CLA methyltransferase (CLAMT). Canonical SLs primarily function as rhizosphere signals, with their biosynthesis in dicots mediated by CYP722C enzymes. It is hypothesized that Arabidopsis does not produce canonical SL because of the lack of the *CYP722C* genes in its genome. Instead, Arabidopsis possesses CYP722A1, a member of the previously uncharacterized CYP722A subfamily, distinct from the CYP722C subfamily.

This study demonstrates that Arabidopsis *cyp722a1* mutants exhibit an earlier floral transition without excessive shoot branching. Biochemical analysis revealed that CYP722A1 catalyzes the hydroxylation of CLA to produce 16-hydroxy-CLA (16-HO-CLA), which is subsequently methylated by CLAMT to form 16-HO-MeCLA. 16-HO-CLA and 16-HO-MeCLA were detected in the wild-type; however, these compounds were absent in *max1-4* mutant, which is deficient in CLA synthesis, and in *cyp722a1* mutant. These findings indicate the presence of CYP722A1-dependent 16-hydroxylation activity of CLA in Arabidopsis. Moreover, they suggest that hydroxylated CLA specifically regulates floral transition, distinct from branching inhibition. By identifying CYP722A1 as a regulator of floral transition, which is the distinct role of the CYP722A subfamily, this work provides insights into the adaptation of SL structures for specialized biological functions in plant development.

## Introduction

Strigolactones (SLs) are a structurally related group of plant apocarotenoids recognized for their diverse roles in plant development and rhizosphere signaling. In the rhizosphere, root-secreted SLs serve as key signals stimulating the germination of root parasitic weeds from the Orobanchaceae family, such as *Striga* spp. and *Orobanche* spp. (Cook et al., 1966; Yoneyama et al., 2010). SLs also promote hyphal branching in arbuscular mycorrhizal fungi, facilitating symbiotic interactions with plants (Akiyama et al., 2005). In addition to their exogenous functions in the rhizosphere, SLs act as endogenous plant hormones, regulating plant architecture by inhibiting shoot branching (Gomez-Roldan et al., 2008; Umehara et al., 2008).

SLs exhibit considerable structural diversity and are broadly categorized into two groups: canonical and noncanonical. Canonical SLs are characterized by a tricyclic lactone ring (ABC-ring) connected to a methylbutenolide ring (D-ring), while noncanonical SLs retain the D-ring but lack the B- or C-ring. Canonical SLs are further divided into two subgroups based on the configuration of their C-rings: orobanchol-type SLs, with α-oriented C-rings, and strigol-type SLs, with β-oriented C-rings (Fig. 1). Despite the structural variations of SLs synthesized in different plant species, the upstream biosynthetic pathways leading to the production of carlactonoic acid (CLA) are highly conserved (Fig. 1). This pathway involves a series of enzymatic reactions initiated with the isomerization and cleavage of β-carotene by DWARF27 (D27), CAROTENOID CELAVAGE DIOXYGENASE 7 (CCD7), and CCD8. This results in the production of carlactone (CL), a key biosynthetic intermediate (Alder et al., 2012; Seto et al., 2014). CL is subsequently oxidized to CLA by cytochrome P450 in the CYP711A subfamily (Abe et al., 2014; Yoneyama et al., 2018; Zhang et al., 2014). Downstream of the CL and CLA, a variety of SL structures are biosynthesized (Nomura et al., 2024).

**Figure 1.**
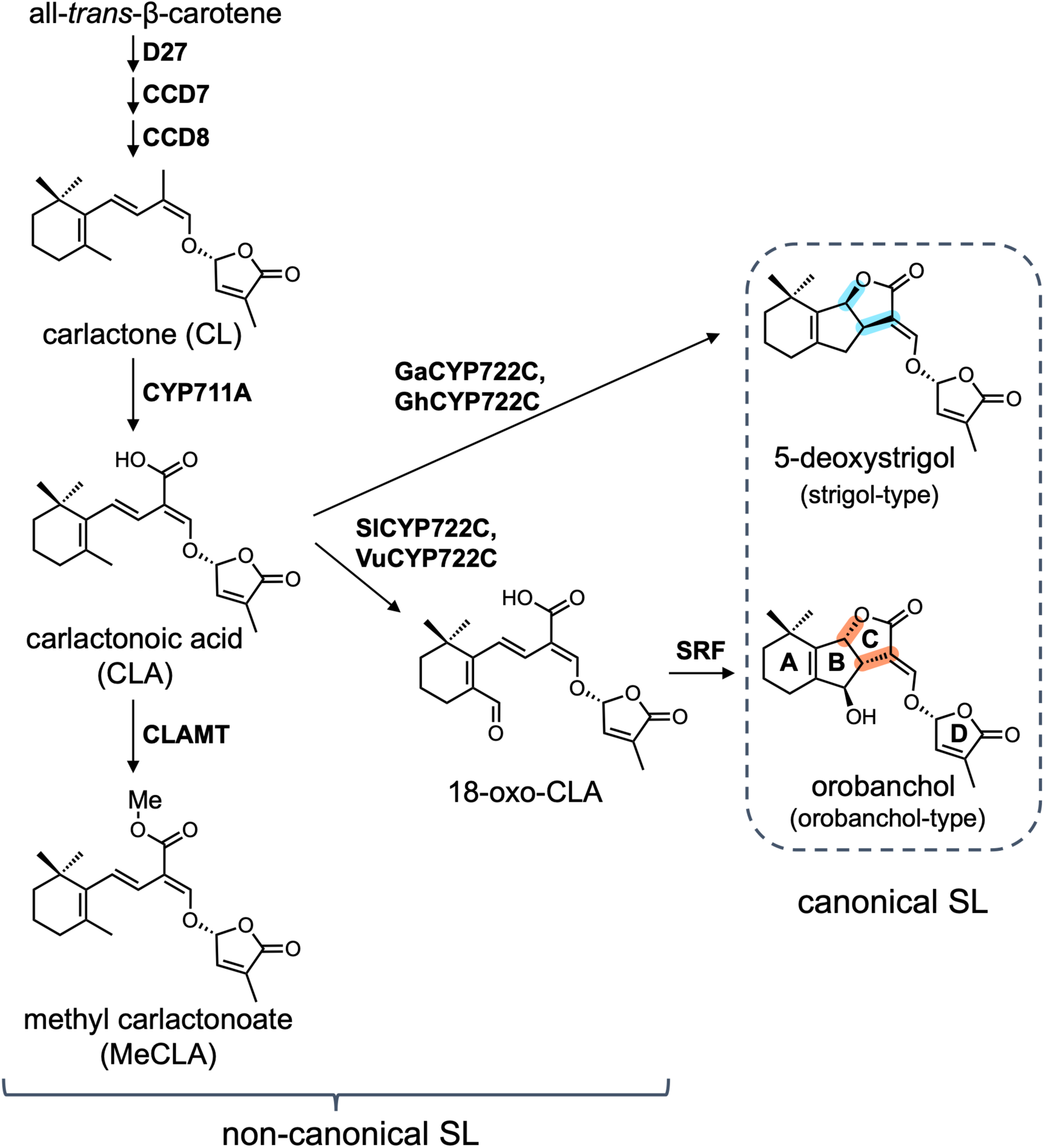
Overview of a part of the strigolactone biosynthesis pathway. The pathway from the biosynthetic intermediate carlactone (CL) to methyl carlactonoate (MeCLA) is proposed to be conserved across many plant species. In canonical SL biosynthesis in dicot plants, CYP722C acts as a key enzyme. In Arabidopsis, canonical SLs are presumed not to be synthesized. Ga, *Gossypium arboreum*; Gh, *Gossypium hirsutum*; Sl, *Solanum lycopersicum*; Vu, *Vigna unguiculata*.

Plants deficient in these biosynthetic genes exhibit a typical SL-deficiency phenotype, most notably increased shoot branching. In addition to shoot branching regulation, SLs influence various plant developmental processes (Bhoi et al., 2021), including floral transition. Recent studies have demonstrated that *Arabidopsis thaliana* mutants with defective SL biosynthesis or signaling pathways exhibit early flowering under both long-day and short-day conditions, suggesting that endogenous SL signaling delay floral transition (Bai et al., 2024; Zhang et al., 2019).

Advances in understanding the downstream biosynthetic pathways of CL and CLA have revealed the key enzymes involved in individual SL production (Nomura et al., 2024). Among these, enzymes from the CYP722C subfamily have been recognized as the drivers of canonical SL biosynthesis in dicots (Homma et al., 2024; Wakabayashi et al., 2023, 2020, 2019) (Fig. 1). Notably, monocots, such as rice (*Oryza sativa*) lack CYP722C genes. In dicot species, like cotton (*Gossypium arboreum* and *G. hirsutum*), which produces 5-deoxystrigol, a strigol-type SL, GaCYP722C and GhCYP722C catalyze the conversion of CLA to 5-deoxystrigol (Wakabayashi et al., 2023, 2020). In tomato (*Solanum lycopersicum*) and cowpea (*Vigna unguiculata*), which produce orobanchol, an orobanchol-type SL, SlCYP722C and VuCYP722C catalyze the conversion of CLA to 18-oxo-CLA, which then cyclizes to form orobanchol through the action of SRF, a member of the dirigent protein family (Homma et al., 2024; Wakabayashi et al., 2019). The identification of the CYP722C subfamily provides insight into the potential for SLs to exhibit distinct functions based on their structural variations, suggesting that the canonical SLs may not have a significant role in shoot branching regulation. Supporting this, *SlCYP722C* knockout mutant in tomato, which is deficient in canonical SLs, does not exhibit increased shoot branching (Wakabayashi et al., 2019).

In Arabidopsis, CLA methyltransferase (CLAMT) has been identified as a crucial enzyme responsible for noncanonical SL biosynthesis, catalyzing the methylation of CLA to form methyl carlactonoate (MeCLA) (Mashiguchi et al., 2022; Wakabayashi et al., 2021) (Fig. 1). Similarly, in other plant species, homologous CLAMT enzymes methylate CLA-type SLs, indicating a conserved enzymatic function in SL biosynthesis (Haider et al., 2023; Li et al., 2024, 2023). The loss of CLAMT in Arabidopsis results in an increased shoot branching phenotype, highlighting its essential role in synthesizing noncanonical SL-related hormones that inhibit shoot branching.

Canonical SL biosynthesis in dicots is mediated by CYP722C; however, it is generally accepted that Arabidopsis does not produce canonical SLs due to the absence of the *CYP722C* gene in its genome. Instead, Arabidopsis possesses CYP722A1 belonging to the CYP722A subfamily. This subfamily is highly conserved in dicot plants. To investigate the potential role of CYP722A enzymes in SL biosynthesis, we have analyzed the function of CYP722A1, encoded by *AT1g19630*, in Arabidopsis. Our study reveals that CYP722A1 catalyzes the hydroxylation of CLA to produce 16-hydroxy-CLA (16-HO-CLA). Additionally, the *cyp722a1* mutants exhibit an earlier floral development phenotype with only minor effects on shoot branching, indicating the hydroxylation of CLA by CYP722A1, which is primarily linked to floral transition rather than branching inhibition.

## Results

### The CYP722A subfamily is conserved in dicot plants

The CYP722 family comprises multiple subfamilies with distinct evolutionary trajectories. In our previous work, we identified the CYP722C subfamily as a key enzyme responsible for the canonical SL biosynthesis from CLA in dicot plants. However, the functions of the other subfamilies within the CYP722 family are still not fully understood. To address this gap, we conducted a phylogenetic analysis of the CYP722 family, which is classified into four subfamilies: CYP722A, CYP722B, CYP722C, and CYP722D (Hansen et al., 2021) (Fig. 2). The analysis across different plant lineages revealed that CYP722A and CYP722C are present in dicot plants, while CYP722B is specific to monocot plants, and CYP722D is restricted to gymnosperms. Notably, the CYP722A, B, and D subfamilies are clustered in the same branch, while the CYP722C subfamily has branched off from this lineage, likely evolving to specialize in the production of canonical SLs in dicot plants. Although the functions of CYP722A, CYP722B, and CYP722D remain largely unexplored, recent findings, published during the preparation of this manuscript, have provided insight into some aspects of CYP722A function (Zhou et al., 2025). Nonetheless, their broad conservation across seed plants implies their essential roles in plant metabolism and development. In Arabidopsis, the genome contains a single gene from the CYP722A subfamily, *CYP722A1* (*AT1g19630*). Given the evolutionary conservation and potential functional relevance, we focused on characterizing this protein in this study.

**Figure 2.**
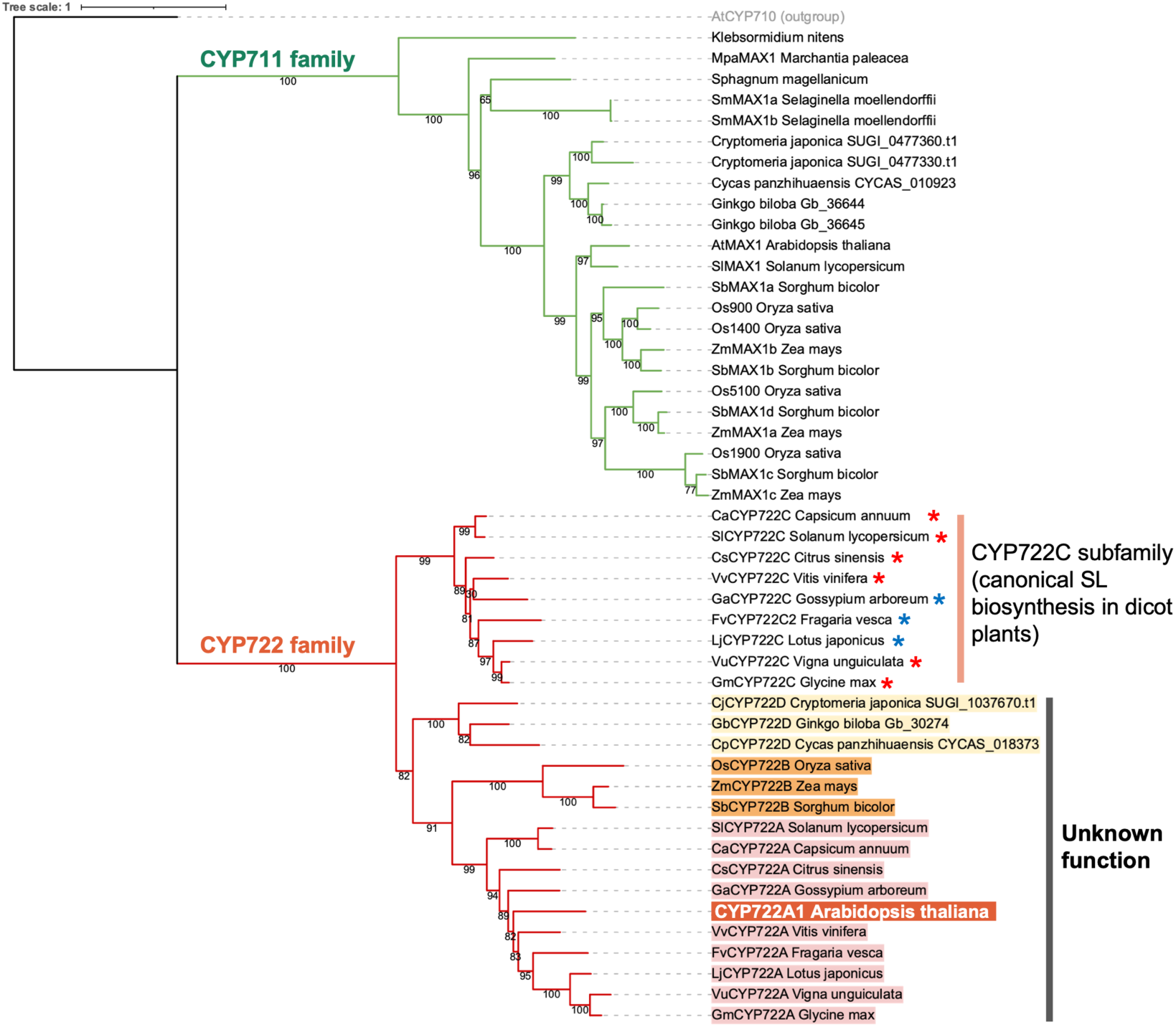
Phylogenetic analysis of SL biosynthesis-related cytochrome P450 enzymes. The analysis was performed using RAxML under the evolutionary model JTT + G + F + I with 900 bootstrap replicates, automatically determined with the option autoMRE. Red asterisks within the CYP722C subfamily indicate proposed involvement in orobanchol biosynthesis, while blue asterisks indicate proposed involvement in 5-deoxystrigol biosynthesis (Wu et al., 2021).

### The *cyp722a1* mutants exhibit an earlier floral transition

To characterize CYP722A1 and investigate its involvement in SL biosynthesis, we analyzed the phenotypes of *cyp722a1* mutant lines (Supplementary Fig. S1). Under long-day conditions with soil-grown plants, *cyp722a1* mutants exhibited increased shoot branching during the early stages of growth (up to approximately 32 days), similar to the SL biosynthesis mutant *max1-4* (*AtCYP711A*-deficient). However, at later growth stages (around 40 days), the number of shoot branches in the *cyp722a1* mutants was comparable to that of the wild-type Col-0 (Fig. 3A and B).

**Figure 3.**
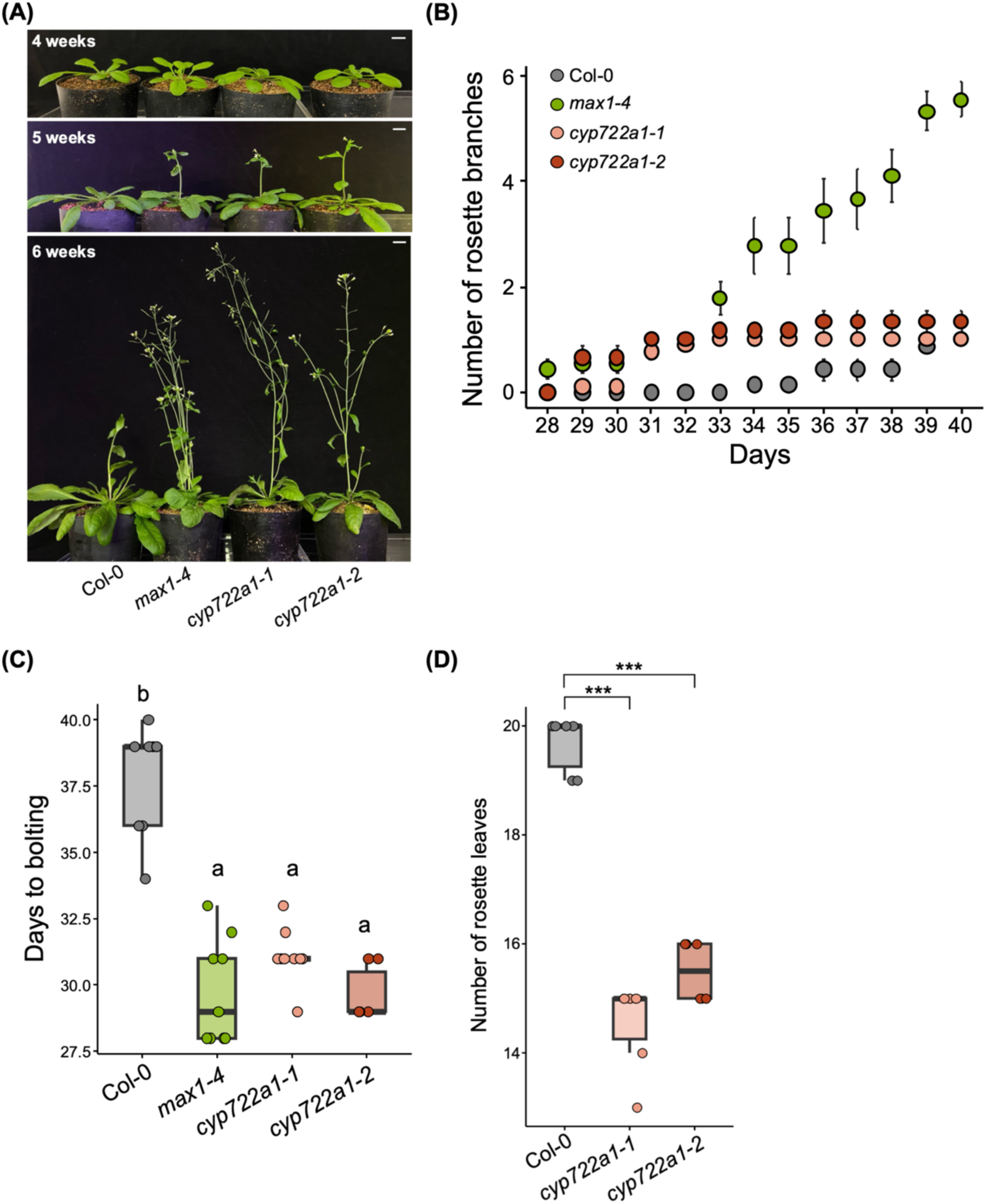
Regulation of floral transition by CYP722A1 in Arabidopsis. (A) Visual phenotypes of flowering time and shoot branching in the SL biosynthesis mutant *max1-4* and *cyp722a1* mutants. Representative images were captured weekly starting from 4 weeks after sowing under long-day conditions. Scale bars = 1 cm. (B) Progression of shoot branching in Arabidopsis WT (Col-0), *max1-4*, and *cyp722a1* mutants. The number of rosette branches (stem length >5 mm) per plant is presented as the mean ± SE. (C) Quantification of the flowering phenotype based on days to bolting. The number of days after sowing until the main stem exceeded 5 mm in length was recorded. Values are shown as individual data points. Different letters indicate significant differences determined by Tukey’s HSD test (*P* < 0.05). In (B) and (C), sample sizes were as follows: Col-0 (*n* = 7), *max1-4* (*n* = 9), *cyp722a1-1* (*n* = 9), and *cyp722a1-2* (*n* = 6). (D) Number of rosette leaves at bolting. The number of rosette leaves was recorded when the main stem exceeded 5 mm in length. Values are shown as individual data points. Significant differences were determined by Dunnett’s test (\*\*\**P* < 0.001, *n* = 6).

Interestingly, both the *max1-4* and *cyp722a1* mutants exhibited an early bolting phenotype (Fig. 3A and C). Moreover, the number of rosette leaves at the bolting stage in the *cyp722a1* mutants was significantly lower than that in Col-0, indicating an accelerated floral transition (Fig. 3D). To verify the association of the mutant phenotype with the gene function, we performed complementation assay by overexpressing *CYP722A1* under CaMV 35S promoter in the *cyp722a1* mutants. Phenotypic analysis of the complemented lines revealed that their bolting phenotypes were partially restored, resulting in delayed bolting compared to the *cyp722a1* mutant (Supplementary Fig. S2A and B). This partial complementation is likely due to the difference in spatial and temporal expression patterns between the 35S and CYP722A promoters. According to the Arabidopsis eFP Browser (Waese et al., 2017), *CYP722A1* is strongly expressed in the first node, second internode, and cauline leaves, suggesting that its precise expression in these tissues is critical for regulating floral transition (Supplementary Fig. S2C). Collectively, these results indicate that *CYP722A1* plays a crucial role in floral transition, similar to other SL biosynthetic genes, but its role in inhibiting shoot branching is relatively minor.

### CYP722A1 catalyzes the conversion of CLA to CLA+16

To investigate the involvement of CYP722A1 in SL biosynthesis, we analyzed its enzymatic activity. Given that members of the CYP722C subfamily catalyze oxygenation reactions with CLA as a substrate (Homma et al., 2024; Wakabayashi et al., 2023, 2020, 2019), we hypothesized that CYP722A1 might also use CLA as a substrate. To test this, we first expressed a recombinant CYP722A1 protein in *Escherichia coli*, truncating its *N*-terminal 26 amino acids, which are predicted to be part of the transmembrane region, and fusing a 6×His-tag to the *C*-terminus (d26CYP722A1). The recombinant d26CYP722A1 was then purified by His-tag affinity chromatography. The expression of active cytochrome P450 enzyme was verified by carbon monoxide (CO) difference spectral analysis, which exhibited a Soret peak with maximum absorbance at 450 nm (Fig. 4A). Using the purified recombinant enzyme, we performed *in vitro* enzyme assays with known endogenous SL precursors of Arabidopsis: CL, CLA, and MeCLA (Fig. 1). Racemic mixtures of CLA and MeCLA (*rac*-CLA and *rac*-MeCLA) were used for the assays. When *rac*-CLA was used as the substrate, a product with a molecular mass 16 Da larger than that of CLA, designated CLA+16, was detected by LC-MS/MS analysis (Fig. 4B). The CLA+16 compound was further analyzed using HR-ESI-MS, and a deprotonated molecule [M − H]^−^ was observed at *m/z* 347.15202. Based on this analysis, its molecular formula was inferred to be C_19_H_24_O_6_, suggesting it to be a hydroxylated form of CLA (calcd. for [C_19_H_24_O_6_ − H]^−^, 347.15001). No hydroxylated products were detected when CL or *rac*-MeCLA was used as the substrates (Supplementary Fig. S3).

**Figure 4.**
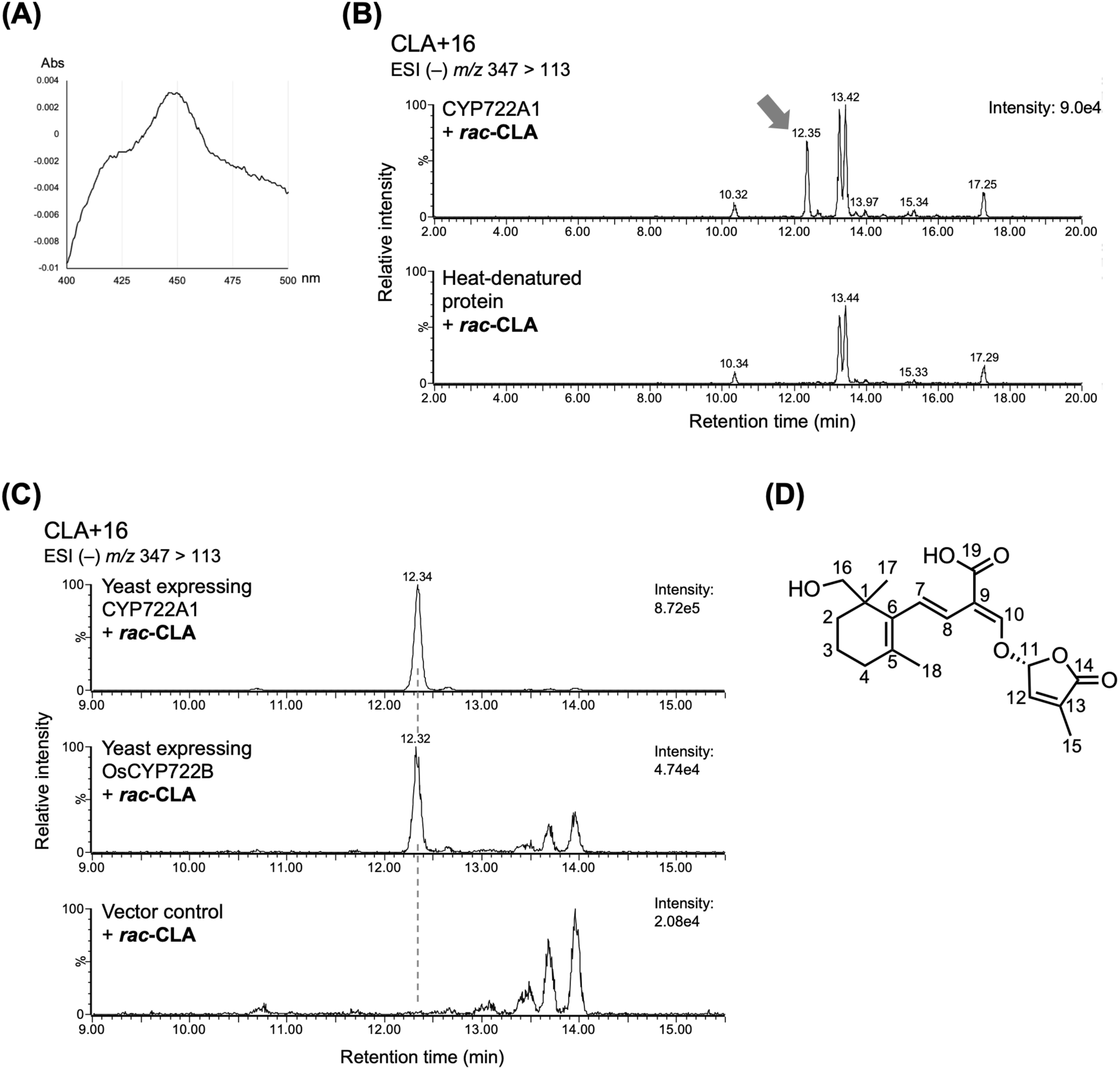
Conversion of CLA to the compound CLA+16 catalyzed by CYP722A1. (A) Carbon monoxide (CO)-difference spectra of the recombinant d26CYP722A1 protein, confirming its expression as in the active form. (B) *In vitro* enzyme assay of recombinant d26CYP722A1. Multiple reaction monitoring (MRM) chromatograms show reaction products when *rac*-CLA is used as a substrate. For the negative control, the assay was performed with heat-denatured recombinant protein. (C) *In vivo* conversion assay using yeast. Yeast expressing CYP722A1 or OsCYP722B was incubated with *rac*-CLA as a substrate, and conversion products were analyzed. Yeast harboring an empty vector served as a negative control. (D) Proposed structure of CLA+16, identified as 16-hydroxy-CLA (16-HO-CLA).

To further validate the enzymatic activity of CYP722A1, we performed a feeding assay using living yeast (*Saccharomyces cerevisiae*) cells (Wu et al., 2021; Yoneyama et al., 2018) as an *in vivo* system. Full-length CYP722A1 was co-expressed with cytochrome P450 reductase in yeast, and the SL precursors were supplied as the substrate for conversion. The reaction products were analyzed by LC-MS/MS. Consistent with the *in vitro* assays, CLA+16 was detected only when *rac*-CLA was used as the substrate (Fig. 4C, Supplementary Fig. S4). Furthermore, a similar assay with OsCYP722B, a rice homolog of CYP722A1, showed that OsCYP722B also catalyzed the conversion of CLA to CLA+16 (Fig. 4C). These results suggest that the CYP722A and CYP722B subfamilies share a conserved enzymatic function in the hydroxylation of CLA to produce CLA+16.

### Structural identification of CLA+16 as 16-hydroxy-CLA, a putative product of CYP722A1

The CLA+16 compound, identified from both the *in vitro* enzyme assay with recombinant protein and the *in vivo* conversion assay with yeast, was analyzed by LC-MS/MS alongside various hydroxylated derivatives of CLA. These derivatives included synthetically prepared 2-HO-CLA, 4-HO-CLA, 5,6-epoxy-CLA, and the enzymatic product of SlCYP722C, 18-HO-CLA (Homma et al., 2024). A comparison of the retention times revealed that the retention time of CLA+16 did not correspond to any of these derivatives (Supplementary Fig. S5), indicating the need for further structural elucidation. To determine the structure of CLA+16, the compound was generated via *in vivo* conversion in yeast expressing CYP722A1. The resulting CLA+16 was extracted from the yeast culture media, partially purified using a solid-phase extraction cartridge, and further purified through reverse-phase HPLC with an ODS column. Finally, CLA+16 was isolated using normal-phase HPLC with a chiral column (Supplementary Fig. S6) and subjected to NMR analysis.

The ^1^H-NMR spectrum of CLA+16 closely resembles that of CLA (Supplementary Table 1 and Supplementary Fig. S7), featuring three methylene signals corresponding to H-2, H-3, and H-4, as well as four methine signals at H-7, H-8, H-10, and H-12. Additionally, three methyl signals were observed for H-15, H-17, and H-18. Notably, the methyl proton signal present in CLA was absent in CLA+16. Instead, two hydroxymethylene protons were observed at *δ* 3.30 and *δ* 3.58 (2H, AB quartet, *J* = 11 Hz). Based on this analysis, we identified the structure of CLA+16 as 16-hydroxy CLA (16-HO-CLA) (Fig. 4D).

A recent study also tested the enzymatic activity of CYP722A1 using an *E. coli*–yeast consortium system, a co-culture model combining bacterial and yeast cells (Zhou et al., 2025). This system confirmed the production of CLA+16 and identified it as 16-HO-CLA. These findings, together with our results, provide further validation of the structural determination and emphasize the robustness of CYP722A1 activity across diverse experimental systems.

### Regulation of floral transition by CLAMT downstream of CYP722A1

The identification of the CYP722A1 enzyme product as 16-HO-CLA suggests that the biosynthetic pathway leading to16-HO-CLA play a critical role in regulating floral transition in Arabidopsis. A previous study showed that methylation of CLA by CLAMT is important for the biosynthesis of the shoot branching-inhibiting hormone, as *clamt* mutants exhibit increased shoot branching (Mashiguchi et al., 2022). Furthermore, another study demonstrated that the SL receptor DWARF14 (AtD14) interacts with MeCLA, but not with CLA *in vitro* (Abe et al., 2014), suggesting that methylation of CLA-type SLs is crucial for receptor binding and subsequent hormonal signaling. To investigate the involvement of CLAMT in the floral transition, we examined the bolting phenotype in *clamt* mutants.

The *clamt-1* (No-0 background) mutants exhibited earlier bolting compared with the wild-type No-0 plants, highlighting the essential role of CLAMT in regulating floral transition (Fig. 5A and B), in addition to its established function in shoot branching (Mashiguchi et al., 2022). To further explore the biochemical function of CLAMT, we performed an enzymatic assay to determine whether CLAMT catalyzes the methylation of 16-HO-CLA. LC-MS/MS analysis of the reaction products revealed a peak corresponding to the methylated form of 16-HO-CLA, detected at the multiple reaction monitoring (MRM) transition at *m/z* 363 > 97. The retention time of this peak matched that of the methylated derivative of 16-HO-CLA (16-HO-MeCLA) produced by the TMS-diazomethane derivatization (Fig. 5C). These findings indicate that CLAMT catalyzes the methylation of 16-HO-CLA to form 16-HO-MeCLA, a result that aligns with the recent report (Zhou et al., 2025). This methylation represents a key step involved in regulating the floral transition downstream of CYP722A1. Furthermore, CYP722A1 did not use MeCLA as a substrate (Supplementary Fig. S3 and S4), suggesting a sequential biosynthetic pathway in which CYP722A1 first hydroxylates CLA to produce 16-HO-CLA, which is then methylated by CLAMT to form 16-HO-MeCLA.

**Figure 5.**
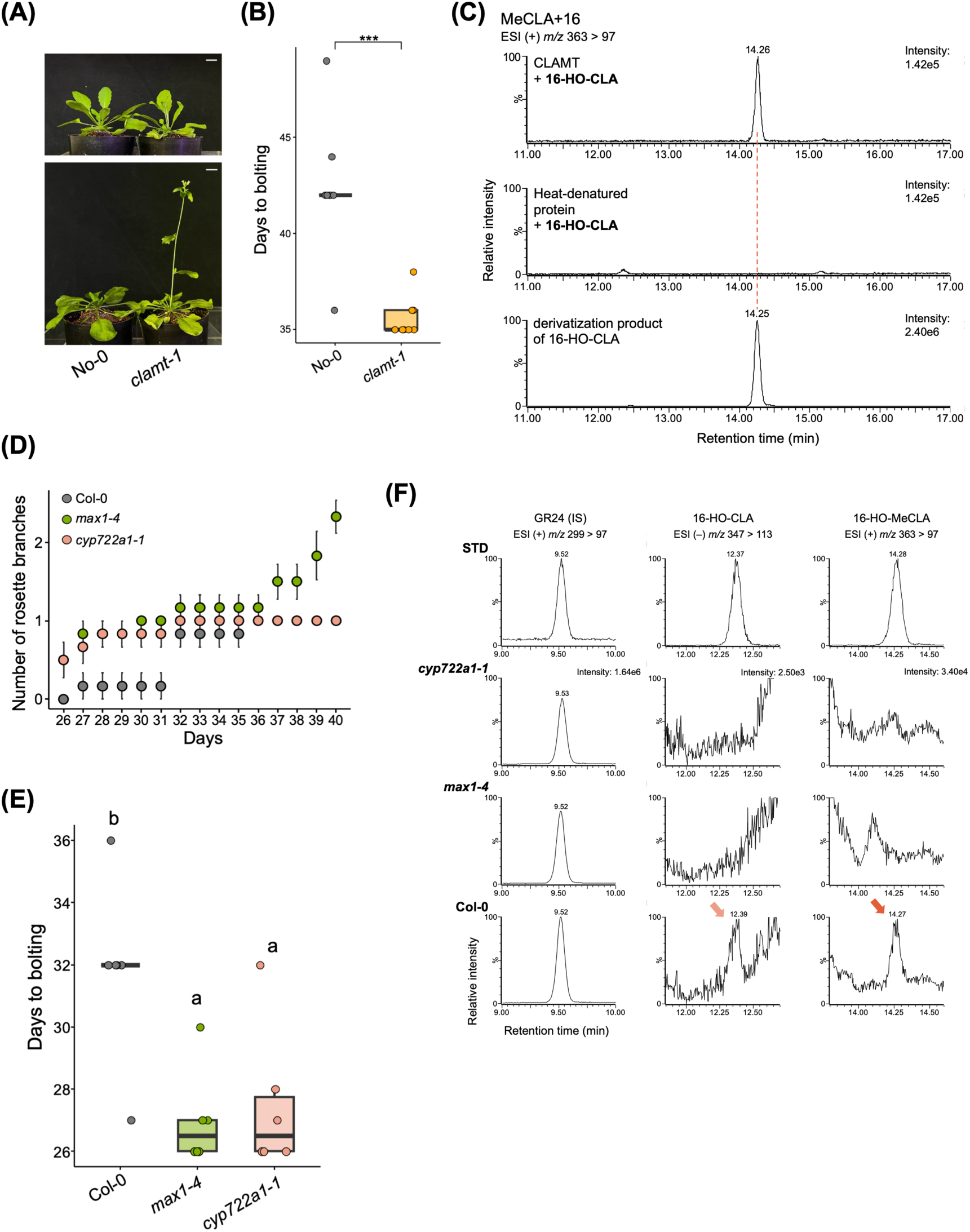
Regulation of floral transition by CLAMT downstream of CYP722A1. (A) Visual phenotypes of flowering time in Arabidopsis WT (No-0) and the SL biosynthesis mutant *clamt-1*. Representative images were captured at 5 and 6 weeks after sowing under long-day conditions. Scale bars = 1 cm. (B) Quantification of the flowering phenotype based on days to bolting. The number of days after sowing until the main stem exceeded 5 mm in length was recorded. Significant differences were determined by Student’s *t-*test (\*\*\**P* < 0.001, No-0, *n* = 9; *clamt-1*, *n* = 9). (C) *In vitro* enzyme assay of recombinant CLAMT. The reaction was performed using 16-HO-CLA, enzymatically produced by CYP722A1, as a substrate. Reaction products were analyzed by LC-MS/MS. For the negative control, the assay was conducted using heat-denatured recombinant protein. The bottom chromatogram shows the methylated derivative of 16-HO-CLA obtained through TMS-diazomethane treatment. (D and E) Phenotypic analysis of Arabidopsis WT (Col-0), *max1-4*, and *cyp722a1-1* mutants under hydroponic conditions. (D) Progression of shoot branching under hydroponic conditions (*n* = 6, mean ± SE). (E) Quantification of the flowering phenotype based on days to bolting. Values are shown as individual data points. Different letters indicate significant differences determined by Tukey’s HSD test (*n* = 6, *P* < 0.05). (F) Analysis of endogenous 16-HO-CLA and 16-HO-MeCLA in Arabidopsis. GR24 was used as internal standard (IS). Arrows indicate distinct peaks in chromatograms corresponding to the retention times of enzymatically produced 16-HO-CLA and 16-HO-MeCLA.

A previous study identified 16-HO-CLA as an endogenous SL in both wild-type Col-0 and the *atd14* mutant. Additionally, 16-HO-MeCLA was detected in the *atd14* mutant (Yoneyama et al., 2020). To further explore the presence of endogenous SLs in Arabidopsis, we analyzed the Col-0, *max1-4*, and *cyp722a1-1* mutants. For the analysis of endogenous SLs, hydroponically grown Arabidopsis plants were used. Even under hydroponic conditions, the *cyp722a1-1* mutant did not exhibit an excessive shoot branching phenotype, and both *max1-4* and *cyp722a1-1* mutants showed earlier bolting phenotypes compared to Col-0 plants (Fig. 5D and E). LC-MS/MS chromatogram of Col-0 extracts showed distinct peaks corresponding to enzymatically produced 16-HO-CLA and 16-HO-MeCLA, thus confirming their presence. In contrast, the chromatograms of the *max1-4* and *cyp722a1-1* mutants displayed significantly lower signal-to-noise ratios at these retention times, with no distinguishable peaks (Fig. 5F). The absence of 16-HO-CLA and 16-HO-MeCLA in the *cyp722a1* mutants was consistent with the recent report (Zhou et al., 2025). These results strongly suggest the absence of these SLs in the mutants and provide additional support for the proposed pathway from CLA to 16-HO-MeCLA. Taken together, these results strongly suggest a relationship between the CYP722A1-mediated biosynthetic pathway from CLA to 16-HO-MeCLA and the regulation of floral transition in Arabidopsis.

## Discussion

In this study, we demonstrated that CYP722A1 plays a crucial role in the regulation of floral transition in Arabidopsis, as evidenced by the earlier bolting phenotype observed in *cyp722a1* mutants. SL biosynthesis mutants typically exhibit characteristic above-ground phenotypes, such as increased shoot branching and earlier floral transition (Bai et al., 2024; Gomez-Roldan et al., 2008; Umehara et al., 2008; Zhang et al., 2019). However, unlike other SL biosynthesis mutants, the *cyp722a1* mutants did not exhibit excessive shoot branching (Fig. 3). This suggests that CYP722A1 mediates a specialized SL biosynthesis pathway primarily involved in floral transition regulation rather than in shoot branching inhibition.

Through *in vitro* enzyme assays with recombinant enzyme and *in vivo* conversion assays in yeast, we demonstrated that CYP722A1 catalyzes the hydroxylation of CLA to produce 16-HO-CLA (Fig. 4). The recent study using the *E. coli*–yeast consortium system similarly reported that CYP722A1 catalyzes the conversion of CLA to 16-HO-CLA (Zhou et al., 2025). While this co-culture system is an innovative and powerful tool for enzyme functional analysis, it requires precise interpretation to ensure accuracy. For example, a prior report using this system suggested that SlCYP722C catalyzes the stereoselective conversion of CLA to orobanchol (Wu et al., 2021). However, we later clarified that SlCYP722C specifically catalyzes the oxidation of CLA to 18-oxo-CLA, with the subsequent conversion to orobanchol requiring the involvement of a distinct enzyme, SRF (Homma et al., 2024). In this study, we utilized both the *in vivo* yeast assay system and *in vitro* assays with recombinant d26CYP722A1 expressed in *E. coli*. These complementary approaches allowed us to systematically validate the enzymatic function of CYP722A1. This approach not only corroborates the findings of the recent study but also reinforces the reliability and robustness of our conclusions.

Further *in vitro* assays confirmed that 16-HO-CLA is methylated by CLAMT to form 16-HO-MeCLA. The detection of 16-HO-CLA and 16-HO-MeCLA as endogenous SLs in wild-type Arabidopsis, along with the absence of detectable levels of these compounds in *max1-4* and *cyp722a1-1* mutants (Fig. 5), underscores the reliance of these compounds on the CYP722A1-mediated biosynthetic pathway. Notably, previous studies reported the detection of 16-HO-CL, a hydroxylated derivative of CL, in *max1* mutants (Yoneyama et al., 2020). However, our results indicate that CYP722A1 does not catalyze the hydroxylation of CL (Supplementary Fig. S3 and S4), suggesting that a different enzyme is responsible for the production of 16-HO-CL, thereby adding complexity to the SL biosynthetic pathways.

Previous research has shown that tomato and rice mutants deficient in canonical SLs exhibit normal shoot branching or tillering phenotypes (Ito et al., 2022; Wakabayashi et al., 2019). These findings indicate that canonical SLs are not the primary regulators of shoot branching; instead, they propose the concept of functional differentiation of SLs based on their structures. Our study aligns with this concept by demonstrating that specific SL structures, such as 16-HO-CLA, 16-HO-MeCLA, or their metabolites, may act as key regulators of floral transition in Arabidopsis. We propose that the SL biosynthetic pathway in Arabidopsis diverges into two distinct branches with specialized functions: one governed by the methylation of CLA by CLAMT, which regulates shoot branching inhibition, and another involving CYP722A1 and CLAMT, which regulates floral transition (Fig. 6). Interestingly, the increased shoot branching observed during the early growth stages of *cyp722a1* mutants (Fig. 3B) suggests a potential role for CYP722A1 in shoot branching regulation. Supporting this idea, the recent study reported that exogenous application of 16-HO-CLA to SL-deficient Arabidopsis mutants inhibited excessive shoot branching (Zhou et al., 2025), suggesting that 16-HO-CLA or its metabolites may play functional roles in shoot branching regulation.

**Figure 6.**
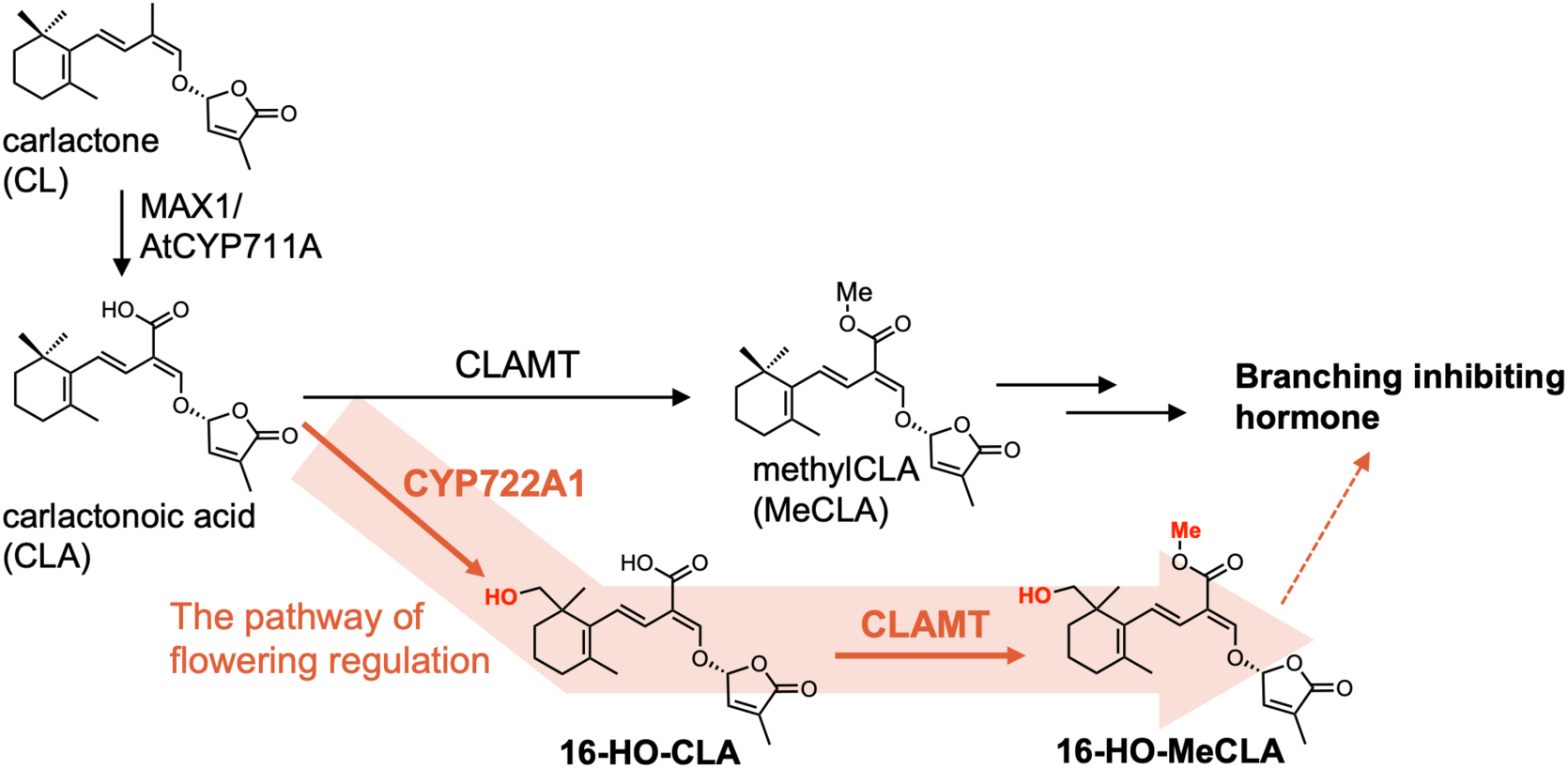
Proposed divergent SL biosynthesis pathway in Arabidopsis. This study proposes a functionally diverged SL biosynthetic pathway in Arabidopsis, with CLA serving as the branching point. One pathway, mediated by CLAMT, synthesizes MeCLA and primarily regulates shoot branching inhibition. The other pathway involves CYP722A1, which converts CLA to 16-HO-CLA, followed by CLAMT-mediated synthesis of 16-HO-MeCLA, primarily regulating floral transition. The CYP722A1-mediated pathway may also partially contribute to branching inhibition.

Our results also highlight the evolutional conservation of CYP722A1 in Arabidopsis and OsCYP722B in rice, both of which catalyze the conversion of CLA to 16-HO-CLA (Fig. 4C). Furthermore, it was recently reported that enzymes belonging to the CYP722A/B subfamilies from other plant species also catalyze similar reactions (Zhou et al., 2025). The CYP722 family first emerged in gymnosperms (Hansen et al., 2021; Vinde et al., 2022) (Fig. 2). The presence of the CYP722D subfamily in gymnosperms, along with the CYP722A and CYP722B subfamilies in angiosperms, suggests that these enzymes collectively contribute to fundamental roles in plant development across seed plants. The SL biosynthesis pathways mediated by the conversion of CLA to 16-HO-CLA appears to be crucial mechanism in plant development, highlighting the potential relevance of these enzymes across diverse plant lineages.

The CYP711A subfamily, which is conserved even in early land plants like liverworts and mosses (Fig. 2), represents one of the earliest cytochrome P450 subfamilies to emerge in SL biosynthesis (Hansen et al., 2021; Kodama et al., 2022; Vinde et al., 2022). Gene duplication within the CYP711A subfamily in monocot grasses has led to the emergence of multiple subfamily members, whereas most other species typically retain only one or two CYP711A genes (Hansen et al., 2021; Yoneyama et al., 2018). This duplication likely facilitated the diversification of SL structures, particularly canonical SL synthesis. For example, in rice, CYP711A2/Os900 and CYP711A3/Os1400 catalyze the conversion of CL to 4-deoxyorobanchol via CLA and the subsequent conversion of 4-deoxyorobanchol to orobanchol, respectively (Yoneyama et al., 2018; Zhang et al., 2014). In sorghum, CYP711A35/SbMAX1a facilitates the conversion of CLA to 5-deoxystrigol, working in concert with other enzymes such as 2-oxoglutarate-dependent dioxygenase and sulfotransferase (Yoda et al., 2023). In dicot plants, the evolutionary expansion of the CYP722 family led to the emergence of the CYP722C subfamily, which facilitates the biosynthesis of canonical SLs. This diversification suggests that dicot plants independently acquired canonical SL biosynthetic capabilities by extending the functional spectrum of the CYP722 family. Further comprehensive functional analyses of the more ancestral subfamilies within the CYP722 family could provide critical insights into the primitive and conserved roles of SLs. One such role, as demonstrated in this study, may involve the regulation of floral development, underscoring the evolutionary significance of these enzymes in plant development.

To gain a more comprehensive understanding of the physiological responses mediated by 16-HO-CLA and 16-HO-MeCLA, further studies are required. These studies should investigate whether the exogenous application of these SLs to mutants can complement their phenotypes and explore their interaction with the SL receptor and downstream signaling pathways. By elucidating the functional diversification of SLs based on their structures, this study provides critical insights that could advance the precise regulation of SL activity and its broader implications in plant development.

## Materials and Methods

### Chemicals

Authentic samples of CL, *rac-*CLA, and *rac*-MeCLA were prepared as previously described (Iseki et al., 2018; Wakabayashi et al., 2022, 2019). Synthesis of 2-HO-CLA, 4-HO-CLA, 5,6-epoxy-CLA is described in the Supplementary Materials and Methods.

### Plant materials and growth conditions

Arabidopsis ecotypes Col-0 and No-0 were used as wild-type plants in this study. The mutants included *max1-4* (Col-0 background) (Abe et al., 2014), *clamt-1* (No-0 background) (Mashiguchi et al., 2022), and *cyp722a1-1* (SALK_088395) and *cyp722a1-2* (SALK_004781), both T-DNA insertion lines in the Col-0 background, obtained from the Arabidopsis Biological Resource Center. To complement the *cyp722a1* phenotype, transgenic plants overexpressing *CYP722A1* were generated. The coding sequence of *CYP722A1* was cloned into the pRI201-AN vector (TaKaRa Bio, Shiga, Japan), and the resulting construct was introduced into *cyp722a1-1* plants via the floral dip method using *Agrobacterium tumefaciens* strain EHA105.

All seeds were surface-sterilized and stratified at 4°C for 3 days after sowing. For soil cultivation, seeds were sown directly onto soil and grown under long-day conditions (16-h-light/8-h-dark photoperiod) at 22°C. For hydroponic cultivation, seeds were germinated on 1/2 MS medium and grown for 1 week before being transferred to containers filled with hydroponic solution (Norén et al., 2004). Hydroponically grown plants were maintained under the same conditions as those used for soil cultivation. The number of rosette leaves and the bolting stage were recorded when plants developed a stem of 5 mm in height.

### Phylogenetic tree analysis

The amino acid sequences used for the phylogenetic analysis were obtained from the following databases: the National Center for Biotechnology Information, Plant GARDEN (Ichihara et al., 2023), ForestGEN (FORest EST and GENome database [https://forestgen.ffpri.go.jp/jp/index.html]), and GinkgoDB (Gu et al., 2022). Accession numbers for the sequences used in this study are provided in Supplementary Data. Cytochrome P450 amino acid sequences were aligned using MAFFT online service (Katoh et al., 2019) and subsequently used for phylogenetic inference. Maximum Likelihood (ML) phylogenetic analysis was performed using RAxML-NG (Kozlov et al., 2019) with the JTT + G + F + I substitution model, identified as optimal based on the Akaike Information Criterion (AIC) determined by ModelTest-NG (Darriba et al., 2020). The number of bootstrap replicates was automatically determined using extended majority rules consensus tree criteria (autoMRE), resulting in 900 replicates.

### *In vitro* enzyme assay using recombinant CYP722A1 expressed in *E. coli*

The expression, purification, and *in vitro* enzyme assay of recombinant CYP722A1 were performed following a previously established method (Homma et al., 2024). Briefly, to produce CYP722A1 in a soluble form, the *N*-terminal 26 residues, predicted to form a transmembrane region, were truncated and replaced with methionine, and a 6×His tag was fused to the *C*-terminus (d26CYP722A1). The modified DNA fragment was cloned into the pColdIII expression vector (TaKaRa Bio, Shiga, Japan) and co-expressed with the chaperone plasmid pGro12 (Nishihara et al., 1998) in *E. coli* BL21 (DE3) cells. Recombinant protein expression was induced, and the protein was extracted and purified using a Ni Sepharose column (Cytiva) according to the previously reported method (Homma et al., 2024). The purified protein was quantified by carbon monoxide (CO)-difference spectral analysis using an extinction coefficient (Σ = 91 mM^−1^ cm^−1^) (Omura and Sato, 1964).

For the *in vitro* enzyme assay, the reaction mixture consisted of 50 mM potassium phosphate buffer (pH 7.4), 50 nM purified recombinant d26CYP722A1, 0.1 unit/mL Arabidopsis NADPH–CYP reductase, 2.5 mM NADPH, and 0.1 μM substrate, in a total volume of 100 μL. Reactions were initiated by adding NADPH and incubated at 30°C for 60 minutes. Reaction products were extracted twice with an equivalent volume of ethyl acetate, and the organic phase was evaporated to dryness. The residues were dissolved in acetonitrile and analyzed by LC-MS/MS.

### *In vivo* conversion using yeast expressing CYP722A1

The *in vivo* conversion assay using living yeast (*S. cerevisiae*) cells was performed by modifying previously reported methods (Wu et al., 2021; Yoneyama et al., 2018). Full-length *CYP722A1* cDNA was cloned into multi-cloning site two of the pELC expression vector, which allows galactose-inducible expression of *Lotus japonicus* CPR1 (LjCPR1) driven by the GAL10 promoter (Seki et al., 2008). The construct was transformed into *S. cerevisiae* INV*Sc*1 (Thermo Fisher Scientific). Recombinant yeast cells were initially cultured in synthetic complete medium without leucine (SC-L) containing 2% (w/v) glucose at 30°C with shaking at 200 rpm for 24 hours.

After the initial culture, cells were collected by centrifugation and resuspended in SC-L medium containing 2% (w/v) galactose to induce protein expression. The cell suspension was adjusted to an OD600 of 0.1, followed by incubation at 30°C with shaking at 200 rpm for 24 hours. The cells were then harvested, resuspended in SC-L medium containing 2% galactose and 10 mM (NH_4_)_2_SO_4_ (Ura et al., 2023) at 1/10 of the original culture volume, and supplemented with the substrate. For the *in vivo* conversion assay, 0.1 μM substrate was used. The cultures were incubated at 22°C with shaking at 200 rpm for 6 hours. Following incubation, yeast cultures were centrifuged, and supernatants were collected to recover *in vivo* conversion products. For extraction, formic acid was added to the supernatant to a final concentration of 0.1%. The mixture was loaded onto an OASIS HLB cartridge (Waters) preconditioned with acetone and equilibrated with water. The cartridge was washed with 40% acetone containing 0.1% formic acid, and the compounds were eluted with acetone containing 0.1% formic acid. The eluted fraction was dried under a stream of nitrogen gas, and the residue was dissolved in acetonitrile for LC-MS/MS analysis.

### *In vitro* enzyme assay of CLAMT

The expression and purification of CLAMT, as well as the enzymatic assay, were performed as previously described (Wakabayashi et al., 2021). The assay was conducted using 16-HO-CLA enzymatically prepared by CYP722A1 as a substrate. Reaction products were analyzed by LC-MS/MS to confirm the catalytic activity of CLAMT.

### Structural determination of CLA+16

Yeast expressing CYP722A1 was cultured in 500 mL of SC-L medium containing 2% galactose. After cultivation, cells were then harvested, resuspended in 50 mL of fresh SC-L medium containing 2% galactose and 10 mM (NH_4_)_2_SO_4_ supplemented with 3 μM *rac*-CLA, and subjected to *in vivo* conversion. The supernatants were collected, and CLA+16 was extracted using the solid-phase extraction method as described above. This process was repeated to accumulate sufficient amounts of CLA+16 for further analysis.

The collected material was further purified by reversed-phase HPLC using a COSMOSIL 5C18-MS-II column (250 × 10 mm i.d., 5 μm) (Nacalai Tesque, Kyoto, Japan). The isocratic system comprised 50 % MeOH/H₂O containing 0.1% formic acid, with a flow rate of 5.0 mL min^−1^. Eluents were monitored at 236 nm, and the peak with a retention time of 68 min was collected (Supplementary Fig. S6A). Subsequently, the compound was purified using normal-phase HPLC on a CHIRALPAK IC column (250 × 10 mm i.d., 5 μm) (Daicel Corporation, Tokyo, Japan). The isocratic system consisted of 30% ethanol/hexane containing 0.1% formic acid, with a flow rate of 1.0 mL min^−1^. Eluents were monitored at 236 nm, and the peak with a retention time of 26 min was collected (Supplementary Fig. S6B).

The purified compound was subjected to NMR analysis. The spectra were recorded in C_6_D_6_ using a Bruker Biospin AC400M spectrometer (400 MHz).

### Extraction of endogenous SLs from Arabidopsis

Endogenous SLs were extracted from Arabidopsis plants (12 plants for a sample) grown hydroponically for 4 weeks. After removing the rosette leaves, the plants were homogenized in acetone and left to stand at 4°C for two days. Following this incubation, GR24 was added as an internal standard (IS) prior to proceeding with the purification steps. The acetone was evaporated under a stream of nitrogen gas, and the residue was resuspended in 10% acetone containing 0.1% formic acid. The suspension was subjected to solid-phase extraction using an OASIS HLB cartridge as described above. The eluted fraction was dried under nitrogen gas, and the residue was re-dissolved in acetonitrile for LC-MS/MS analysis.

### Analysis of SLs

The SLs were analyzed using an LC-MS/MS system (Waters) consisting of an Acquity UPLC and a Xevo TQ-S Cronos tandem mass spectrometer. The analytical conditions were as follows: the capillary voltage was set to 0.8 kV for positive ion mode and 0.6 kV for negative ion mode. The source temperature was maintained at 120°C, and the desolvation gas temperature was set to 500°C. The cone and desolvation N_2_ gas flow rates were 50 and 1000 L h^−1^, respectively. Chromatographic separation was performed with an ODS column (COSMOSIL 2.5C18-MS-II, 100 × 2.0 mm i.d., 2.5 µm; Nacalai Tesque, Kyoto, Japan) at a column oven temperature of 40°C. The elution was performed in a linear gradient system using solvent A (H₂O with 0.1% formic acid) and solvent B (MeOH with 0.1% formic acid) under the following conditions: 0–0.75 min, 10% B; 0.75–1.5 min, 10–35% B; and 1.5–24 min, 35–95% B, at a flow rate of 0.3 mL min⁻¹. Multiple reaction monitoring (MRM) was used to detect SLs, and the MRM transitions were set based on previously reported conditions (Yoneyama et al., 2020). The transitions were as follows: CL, *m/z* 303.2 > 97.0 (positive mode); CL+16, *m/z* 301.2 > 97.0 (positive mode); CLA, *m/z* 331.2 > 113.0 (negative mode); CLA+16, *m/z* 347.2 > 113.0 (negative mode); MeCLA, *m/z* 347.2 > 97.0 (positive mode); and MeCLA+16, *m/z* 363.2 > 97.0 (positive mode). Data acquisition and analysis were performed using MassLynx 4.2 software (Waters).

## Data Availability Statement

The data supporting the findings of this study are available within the article, and Supplementary Data.

## Funding

This work was supported by Japan Science and Technology Agency (JST) PRESTO (No. JPMJPR22DA to T.W.), by Japan Society for the Promotion of Science (JSPS) KAKENHI (No. 20K15459 to T.W.).

## Acknowledgements

We are grateful to Dr. Toshiya Muranaka and Dr. Hikaru Seki at Osaka University for providing the pELC expression vector. The authors would like to thank Enago (www.enago.jp) for the English language review.

## Author Contributions

Conceptualization: T. Wakabayashi and Y. Sugimoto. Methodology: H. Takikawa, T. Wakabayashi, M. Kuno, and M. Homma. Investigation: M. Kuno, A. Miyamoto, H. Takano, M. Homma, N, Shiotani, K. Uchida, and T. Wakabayashi. Project administration: H. Takikawa, M. Mizutani, and Y. Sugimoto. Writing – original draft: T. Wakabayashi. Writing–review and editing: H. Takikawa, M. Nakajima, M. Mizutani, T. Wakabayashi, and Y. Sugimoto.

## Disclosures

Conflicts of interest: No conflicts of interest declared.

## Notes

### Competing Interest Statement

The authors have declared no competing interest.

