## Supplementary Data for "CYP722A1-driven 16-hydroxylation of carlactonoic acid regulates the floral transition in Arabidopsis"

**Supplementary Materials and Methods**

**Synthesis of CLA derivatives**

4-HO-CLA, 5,6-epoxy-CLA, and 2-HO-CLA were synthesized using the following procedures.

*Synthesis of 4-HO-CLA*

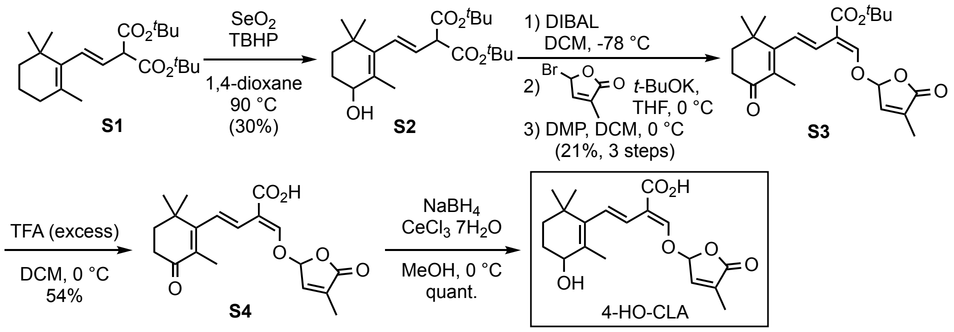


*Synthesis of 5,6-epoxy-CLA*

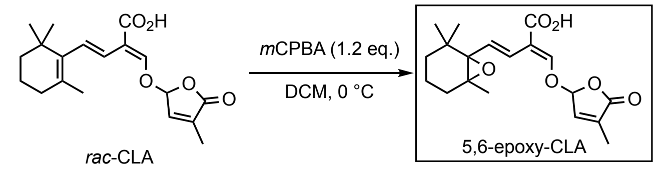


*Synthesis of 2-HO-CLA*

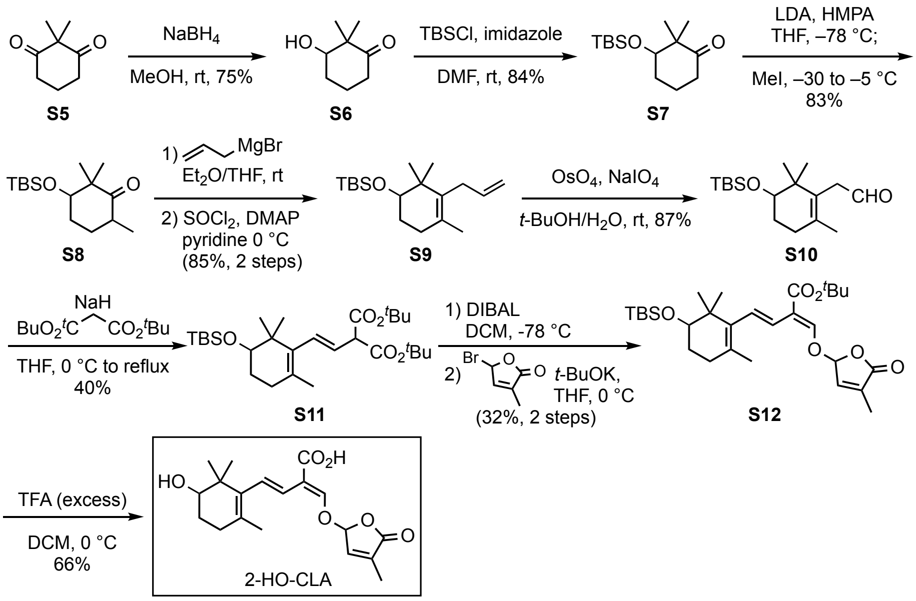


<u>di-*tert*-butyl 2-(2-(3-hydroxy-2,6,6-trimethylcyclohex-1-enyl)vinyl)malonate (**S2**)</u>

To a solution of **S1** (688 mg, 1.88 mmol) and $SeO_2$ (21 mg, 19 mmol) in 1,4-dioxane (18 mL), TBHP 70% in water; 0.26 mL, 1.9 mmol) were added. After stirring at 90 °C for 20 h, the reaction mixture was cooled, diluted with water, and extracted with EtOAc. The organic layer was washed with water and brine, dried with $MgSO_4$, and concentrated under reduced pressure. The residue was purified by $SiO_2$ column chromatography to give **S2** (214 mg, 30%) as a racemic and diastereomeric mixture. Note that the over-oxidized ketone (41 mg, 6%), and the recovered **S1** (166mg, 24%) were also obtained.

$^1$H NMR (400 MHz, $CDCl_3$) δ: 0.97 (3H, s), 1.01 (3H, s), 1.42-1.93 (4H, m), 1.46 (18H, s), 1.82 (3H, br. s), 3.86 (1H, br. d, *J* = 8.8 Hz), 3.97 (1H, m), 5.61 (1H, dd, *J* = 16.0, 8.8 Hz), 6.02 (1H, br. d, *J* = 16.0 Hz).

*tert*-butyl 4-oxocarlactonoate (**S3**)

DIBAL (1.03 M in hexane, 135 µL, 140 µmol) was added to a solution of **S2** (17.7 mg, 46.5 µmol) in DCM (0.5 mL). After stirring at –78 °C for 1 h, the reaction mixture was quenched with MeOH and diluted with sat. Rochelle's salt aq. The resulting mixture was allowed to warm to room temperature with stirring and extracted with DCM. The organic layer was washed with sat. Rochelle's salt aq. and brine, dried with $MgSO_4$, and concentrated under reduced pressure. The residue was used in the next reaction without purification.

*t*-BuOK (5.4 mg, 47 µmol) was added to a solution of the above product (~46.5 µmol) in THF (0.5 mL) at 0 °C. After stirring for 10 min, a solution of 5-bromo-3-methyl-2(5*H*)-furanone (8.2 mg, 47 µmol) in THF (0.15 mL) was added to this solution. After stirring at room temperature for 4 h, the reaction mixture was diluted with $H_2O$ and extracted with DCM. The organic layer was dried with $MgSO_4$ and concentrated under reduced pressure. The residue was diluted with DCM (1 mL), which was added Dess-Martin periodinane (22 mg, 52 µmol). After stirring for 20 min, the reaction mixture was diluted with a mixture of 5% $Na_2S_2O_3$ aq. and sat. $NaHCO_3$ aq. (1:1), and extracted with DCM. The organic layer was washed with a mixture of 5% $Na_2S_2O_3$ aq. and sat. $NaHCO_3$ aq. (1:1), dried with $MgSO_4$, and concentrated under reduced pressure. The residue was purified by $SiO_2$ column chromatography to give **S3** (3.9 mg, 21% in 2 steps).

$^1$H NMR (400 MHz, $C_6D_6$) δ:0.91 (3H, s), 0.95 (3H, s), 1.29 (3H, br. s), 1.39 (9H, s), 1.39 (2H, br. t, *J* = 6.8 Hz), 2.16 (3H, br. s), 2.30 (2H, br. t, *J* = 6.8 Hz), 4.98 (1H, m), 5.64 (1H, m), 6.62 (1H, d, *J* = 16.8 Hz), 7.23 (1H, br. d, *J* = 16.8 Hz), 7.54 (1H, br. s).

4-oxocarlactonoic acid (**S4**)

TFA (0.36 mL, 4.7 mmol) was added to an ice-cooled solution of **S3** (18.9 mg, 47.0 µmol) in DCM (1.0 mL) at 0 °C. After stirring overnight, the reaction mixture was diluted with water and extracted with DCM. The organic layer was washed with water and brine, dried with $MgSO_4$, and concentrated under reduced pressure. The residue was purified by $SiO_2$ column chromatography to give **S4** (8.8 mg, 54%).

$^1$H NMR (400 MHz, $C_6D_6$) δ:0.88 (3H, s), 0.92 (3H, s), 1.28 (3H, br. s), 1.39 (2H, br. t, *J* = 6.8 Hz), 2.13 (3H, br. s), 2.30 (2H, br. t, *J* = 6.8 Hz), 4.93 (1H, br. s), 5.60 (1H, br. s), 6.53 (1H, d, *J* = 16.8 Hz), 7.20 (1H, br. d, *J* = 16.8 Hz), 7.59 (1H, br. s).

4-hydroxycarlactonoic acid (4-HO-CLA)

To a solution of **S4** (1.6 mg, 4.6 μmol) in MeOH (50 μL), $CeCl_3$•$7H_2O$ (7.6 mg, 20 μmol) and $NaBH_4$ (0.4 mg, 10 μmol) were added at 0 °C. After stirring for 25 min, the reaction mixture was diluted with sat. $NH_4Cl$ aq. and extracted with EtOAc. The organic layer was washed with water and brine, dried with $MgSO_4$, and concentrated under reduced pressure to give crude 4-HO-CLA (quant.) as a racemic and diastereomeric mixture. This was employed as an authentic sample without purification.

$^1$H NMR (400 MHz, $C_6D_6$) δ: 0.98, 1.01, 1.04 and 1.07 (total 6H, each s), 1.27 (3H, br. s), 1.91 and 1.93 (total 3H, each br. s), 3.74 (1H, m), 4.94 (1H, m), 5.60 (1H, m), 6.51 (1H, d, *J* = 16.4 Hz), 7.20 (1H, d, *J* = 16.4 Hz), 7.54 (1H, m). (Only clearly observed signals were described.)

5,6-epoxycarlactonoic acid (5,6-epoxy-CLA)

*m*-CPBA (65%; 3.1 mg, 12 μmol) was added to an ice-cooled solution of CLA (3.2 mg, 9.6 μmol) in DCM (0.2 mL) at 0 °C. After stirring for 20 min, the reaction mixture was diluted with 5% $Na_2S_2O_3$ aq. and extracted with DCM. The organic layer was washed with 5% $Na_2S_2O_3$ aq., dried with $MgSO_4$, and concentrated under reduced pressure. The residue was purified by $SiO_2$ column chromatography to give 5,6-epoxyCLA (containing impurities; 1.5 mg, ~50%) as a racemic and diastereomeric mixture. Note that further purification of 5,6-epoxyCLA with $SiO_2$ column chromatography resulted in its decomposition.

$^1$H NMR (400 MHz, $C_6D_6$) δ: 4.89 (1H, br. s), 5.45 (1H, br. s), 6.89 (1H, each br. d, *J* = 16.0 Hz), 6.94 (1H, br. s), 7.36 (1H, br. s), 7.54 (1H, br. s). (Only clearly observed signals were described.)

3-hydroxy-2,2-dimethyl-1-cyclohexanone (**S6**)

$NaBH_4$ (>95%, 117 mg, 2.95 mmol) was added to a solution of **S5** (1.82 g, 13.0 mmol) in MeOH (40 mL) at 0 °C. After stirring at room temperature for 1 h, the reaction mixture was quenched with water and concentrated under reduced pressure to remove MeOH. The resulting mixture was extracted with EtOAc. The organic layer was dried with $Na_2SO_4$ and concentrated under reduced pressure. The residue was purified by $SiO_2$ column chromatography to give **S6** (1.37 g, 75%) as a colorless oil.

$^1$H NMR (400 MHz, $CDCl_3$): δ 1.14 (3H, s), 1.18 (3H, s), 1.68 (1H, m), 1.83 (1H, m), 2.04 (2H, m), 2.40 (2H, m), 3.72 (1H, m).

3-(*tert*-butyldimethylsilyloxy)-2,2-dimethyl-1-cyclohexanone (**S7**)

Imidazole (1.97 g, 28.9 mmol) and TBSCl (1.98 g, 13.1 mmol) were added to a solution of **S6** (1.37 g, 9.67 mmol) in DMF (40 mL). After stirring at 40 °C for 50 h, the reaction mixture was quenched with MeOH, diluted with water, and extracted with $Et_2O$. The organic layer was washed with brine, dried with $Na_2SO_4$, and concentrated under reduced pressure. The residue was purified by $SiO_2$ column chromatography to give **S7** (2.09 g, 84%) as a colorless oil.

$^1$H NMR (400 MHz, $CDCl_3$): δ 0.04 (6H, s), 0.88 (9H, s), 1.08 (3H, s), 1.12 (3H, s), 1.63 (1H, m), 1.99

(1H, m), 2.38 (2H, m), 3.67 (1H, dd, $J$ = 6.8, 2.8 Hz).

3-(*tert*-butyldimethylsilyloxy)-2,2,6-trimethyl-1-cyclohexanone (**S8**)

*n*-BuLi (2.6 M in hexane, 4.1 mL, 11 mmol) was slowly added to a solution of *i*-$Pr_2NH$ (1.4 mL, 10 mmol) in THF at –60 °C. After stirring at –60 °C for 15 min, HMPA (3.0 mL, 17 mmol) was added to the solution. This mixture was then cooled to –78 °C and stirred for 40 min. Then a solution of **S7** (2.07 g, 8.08 mmol) in THF (10 mL) was added to the above prepared LDA solution. After stirring at –78 °C for 35 min, MeI (0.67 mL, 11 mmol) was added to this solution. The reaction mixture was stirred at –30 °C for 1 h and allowed to warm to –5 °C with stirring for 1.5 h. The reaction mixture was quenched with sat. $NH_4Cl$ aq. and extracted with $Et_2O$. The organic layer was washed with brine, dried with $Na_2SO_4$, and concentrated under reduced pressure. The residue was purified by $SiO_2$ column chromatography to give **S8** (1.81 g, 83%) as a diastereomer mixture.

$^1$H NMR (400 MHz, $CDCl_3$): δ 0.02–0.06 (6H, m), 0.84–0.90 (9H, m), 0.97–1.01 (3H, m), 1.05–1.09 (3H, m), 1.57 (3H, s), 1.66 (1H, m), 1.76–1.93 (2H, m), 2.16 (1H, m), 2.63 (1H, m), 3.83 (1H, brs).

2-allyl-4-(*tert*-butyldimethylsilyloxy)-1,3,3-trimethyl-1-cyclohexene (**S9**)

To a mixture of Mg (0.75 g, 31 mmol) and $I_2$ (a small piece) in $Et_2O$ (30 mL) was added allyl bromide (2.8 mL, 32 mmol) slowly at 0 °C. After stirring at room temperature for 45 min, a solution of **S8** (0.87 g, 3.2 mmol) in THF (8 mL) was slowly added at 0 °C. After stirring at room temperature for 2 h, the reaction mixture was quenched with sat. $NH_4Cl$ aq. and extracted with $Et_2O$. The organic layer was dried with $Na_2SO_4$, and concentrated under reduced pressure to give a yellow oil, which was used in the next reaction without further purification.

$SOCl_2$ (0.30 mL, 4.1 mmol) was added to an ice cooled solution of the above product (~3.2 mmol) and DMAP (105 mg, 0.859 mmol) in pyridine (16 mL) at 0 °C. After stirring at 0 °C for 1.5 h, the reaction mixture was quenched with water and extracted with EtOAc. The combined organic layer was washed with brine, dried with $Na_2SO_4$, and concentrated under reduced pressure. The residue was purified by $SiO_2$ column chromatography to give **S9** (0.80 g, 85% in 2 steps).

$^1$H NMR (400 MHz, $CDCl_3$): δ 0.03–0.05 (6H, m), 0.89 (9H, s), 0.91 (3H, s), 0.98 (3H, s), 1.55 (3H, s), 1.62–1.67 (2H, m), 2.01–2.04 (2H, m), 2.77 (1H, t, $J$ = 5.6 Hz), 3.47 (1H, t, $J$ = 6.4 Hz), 4.49–4.98 (2H, m), 5.75 (1H, ddd, $J$ = 16.4, 10.8, 6.4 Hz).

2-(5-(*tert*-butyldimethylsilyloxy)-2,6,6-trimethylcylohex-1-enyl)acetaldehyde (**S10**)

$NaIO_4$ (387 mg, 1.81 mmol) and $OsO_4$ (20 mg/mL in *t*-BuOH, 0.30 mL, 24 μmol) were added to a solution of **S9** (132 mg, 0.446 mmol) in *t*-BuOH/$H_2O$ (v/v, 3:1, 4 mL). After stirring at room temperature for 3 h, the reaction mixture was quenched with sat. $Na_2SO_3$ aq. and filtered through Celite. The filtrate was diluted with $H_2O$ and extracted with EtOAc. The organic layer was washed with sat. $Na_2SO_3$ aq. and brine, and concentrated under reduced pressure. The residue was purified by $SiO_2$ column chromatography to give **S10** (0.1147 g, 87%).

$^1$H NMR (400 MHz, $CDCl_3$): δ 0.03–0.06 (6H, m), 0.89 (9H, s), 0.90 (3H, s), 0.96 (3H, s), 1.56 (3H, s), 1.65–1.70 (2H, m), 2.08–2.11 (2H, m), 3.09 (2H, s), 3.52 (1H, t, *J* = 6.8 Hz), 9.51 (1H, t, *J* = 2.4 Hz).

di-*tert*-butyl (*E*)-2-(2-(5-(*tert*-butyldimethylsilyloxy)-2,6,6-trimethylcyclohex-1-enyl)vinyl)malonate (**S11**)

Di-*tert*-butyl malonate (2.2 mL, 9.9 mmol) was added to a suspension of NaH (55% dispersion, 0.45 g, 10 mmol) in THF (40 mL). After stirring at 0 °C for 45 min, a solution of **S10** (1.43 g, 4.83 mmol) in THF (10 mL) was added to this mixture at room temperature. After stirring under reflux for 15 h, the reaction mixture was cooled to room temperature, quenched with sat. $NH_4Cl$ aq., and extracted with EtOAc. The organic layer was washed with brine, dried with $Na_2SO_4$, and concentrated under reduced pressure. The residue was purified by $SiO_2$ column chromatography to give **S11** (0.962 g, 40%) as a pale yellow oil.

$^1$H NMR (400 MHz, $CDCl_3$): δ 0.02–0.07 (6H, m), 0.89 (9H, s), 0.93 (3H, s), 0.96 (3H, s), 1.46 (18H, s), 1.55 (3H, s), 1.63–1.69 (2H, m), 2.03–2.08 (2H, m), 3.47 (1H, dd, *J* = 8.7, 5.2 Hz), 3.87 (1H, d, *J* = 8.7 Hz), 5.54 (1H, dd, *J* = 15.8, 8.7 Hz), 5.99 (1H, d, *J* = 15.8 Hz).

*tert*-butyl 2-(*tert*-butyldimethylsilyloxy)carlactonoate (**S12**)

DIBAL (1.02 M in hexane, 5.0 mL, 5.1 mmol) was added to a solution of **S11** (833 mg, 1.68 mmol) in DCM (8 mL). After stirring at –78 °C for 1 h, the reaction mixture was quenched with MeOH and diluted with sat. Rochelle's salt aq. and EtOAc. The resulting mixture was allowed to warm to room temperature with stirring for 2 h and extracted with EtOAc. The organic layer was washed with brine, dried with $Na_2SO_4$, and concentrated under reduced pressure. The residue was purified by $SiO_2$ column chromatography to give a yellow oil, which was used in the next reaction without further purification.

*t*-BuOK (198 mg, 1.76 mmol) was added to a solution of the above product (~1.68 mmol) in THF (10 mL) at 0 °C. After stirring at room temperature for 50 min, a solution of 5-bromo-3-methyl-2(5*H*)-furanone (308 mg, 1.74 mmol) in THF (4 mL) was added to this solution. After stirring at room temperature for 14 h, the reaction mixture was quenched with sat. $NH_4Cl$ aq. and extracted with EtOAc. The organic layer was dried with $Na_2SO_4$ and concentrated under reduced pressure. The residue was purified by $SiO_2$ column chromatography to give **S12** (278 mg, 32% in 2 steps) as a yellow oil.

$^1$H NMR (400 MHz, $CDCl_3$): δ 0.04–0.07 (6H, m), 0.89 (9H, s), 0.94 (3H, s), 0.98 (3H, s), 1.51 (9H, s), 1.56 (3H, s), 1.63–1.71 (2H, m), 2.01–2.02 (2H, m), 2.05 (3H, s), 3.46 (1H, m), 6.02 (1H, d, *J* = 16.8 Hz), 6.12 (1H, t, *J* = 1.6 Hz), 6.65 (1H, d, *J* = 16.8 Hz), 6.94 (1H, t, *J* = 1.6 Hz), 7.39 (1H, d, *J* = 2.0 Hz).

2-hydroxycarlactonoic acid (2-HO-CLA)

TFA (0.50 mL, 6.5 mmol) was added to a solution of **S12** (278 mg, 0.536 mmol) in DCM (3 mL) at 0 °C.

After stirring at 0 °C for 1.5 h, the reaction mixture was diluted with water and then extracted with DCM. The organic layer was dried with $Na_2SO_4$ and concentrated under reduced pressure. The residue was purified by $SiO_2$ column chromatography to give 2-HO-CLA (123 mg, 66%) as a racemic and diastereomeric mixture.

$^1$H NMR (400 MHz, $CDCl_3$) δ: 1.01 (3H, s), 1.05 (3H, s), 1.73–1.86 (2H, m),1.70 (3H, s), 2.02 (3H, s), 2.05–2.18 (2H, m), 3.55 (1H, dd, *J* = 9.0, 3.0 Hz), 6.07 (1H, d, *J* = 17.4 Hz), 6.17 (1H, brs), 6.76 (1H, d, *J* = 17.4 Hz), 6.96 (1H, s), 7.64 (1H, s). (OH and $CO_2H$ proton was not clearly observed.)

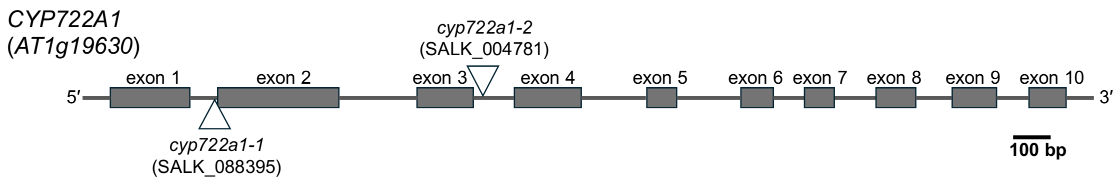


**Supplementary Fig. S1.** Schematic diagram showing the *cyp722a1* mutations.

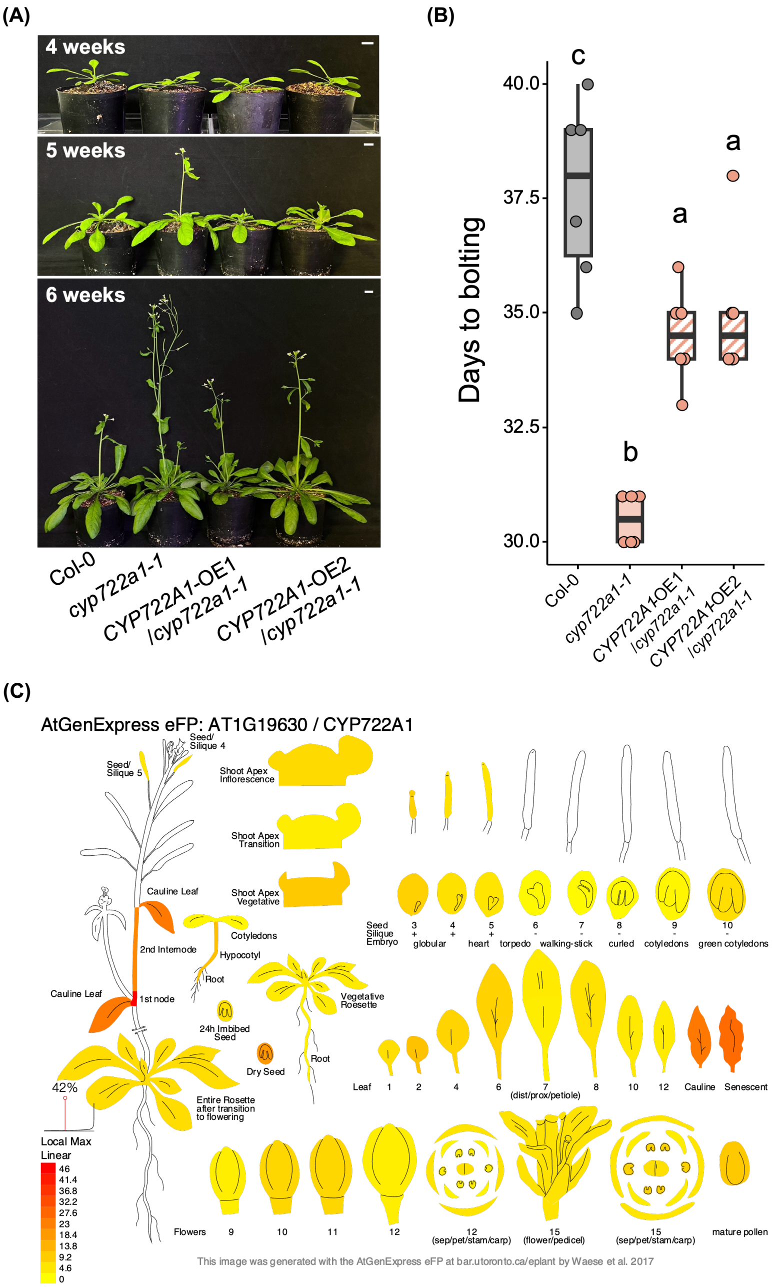


**Supplementary Fig. S2.** Restoration of the early flowering phenotype in the *cyp722a1* mutants by *CYP722A1* overexpression. (A) Visual phenotypes of flowering time in the *cyp722a1* mutants and two independent *CYP722A1* overexpression lines (*CYP722A1*-OE1 and *CYP722A1*-OE2). Representative images were captured weekly starting from 4 weeks after sowing under long-day conditions. Scale bars = 1 cm. (B) Quantification of the flowering phenotype based on days to bolting. The number of days after sowing until the main stem exceeded 5 mm in length was recorded. Values are shown as individual data points. Different letters indicate significant differences determined by Tukey's HSD test ($n = 6$, $P < 0.05$). (C) Digital expression pattern of *CYP722A1* in Arabidopsis tissues visualized using the Arabidopsis eFP browser.

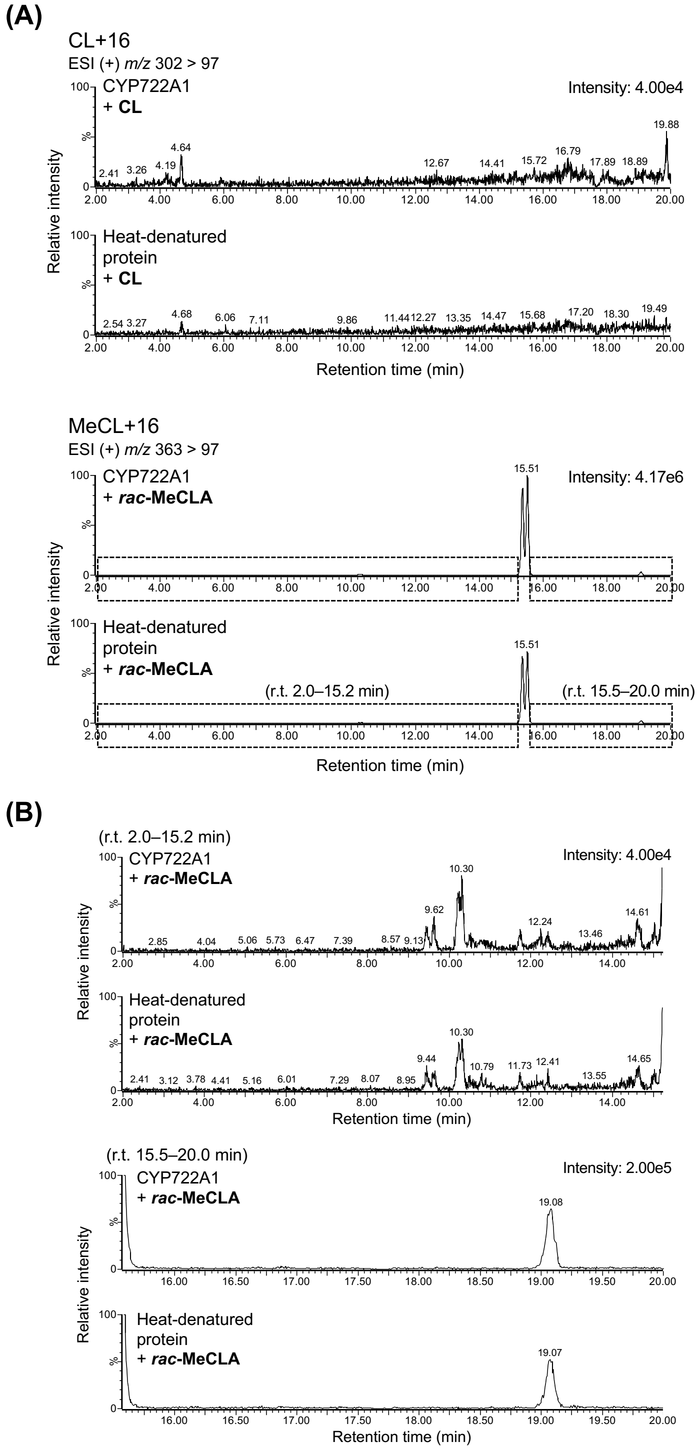


**Supplementary Fig. S3.** Analysis of reaction products in the *in vitro* enzyme assay of CYP722A1 using CL and *rac*-MeCLA as the substrates. (A) MRM chromatograms of hydroxylated products in the reaction mixtures when CL and *rac*-MeCLA were used as the substrates for CYP722A1. (B) Enlarged view of the chromatogram for *rac*-MeCLA.

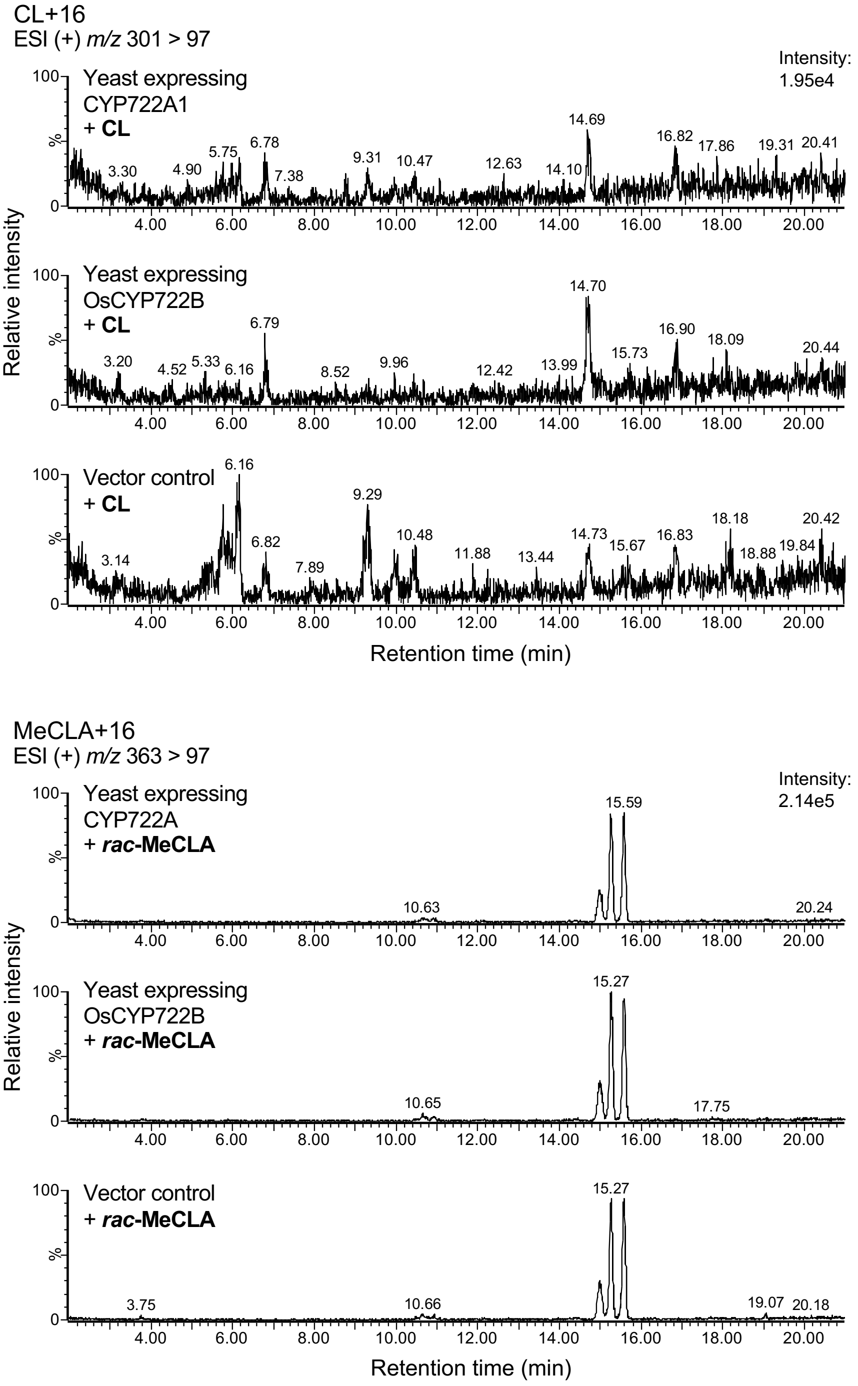


**Supplementary Fig. S4.** MRM chromatograms of hydroxylated products in the culture medium from the *in vivo* conversion assays using living yeast cells fed with CL and *rac*-MeCLA.

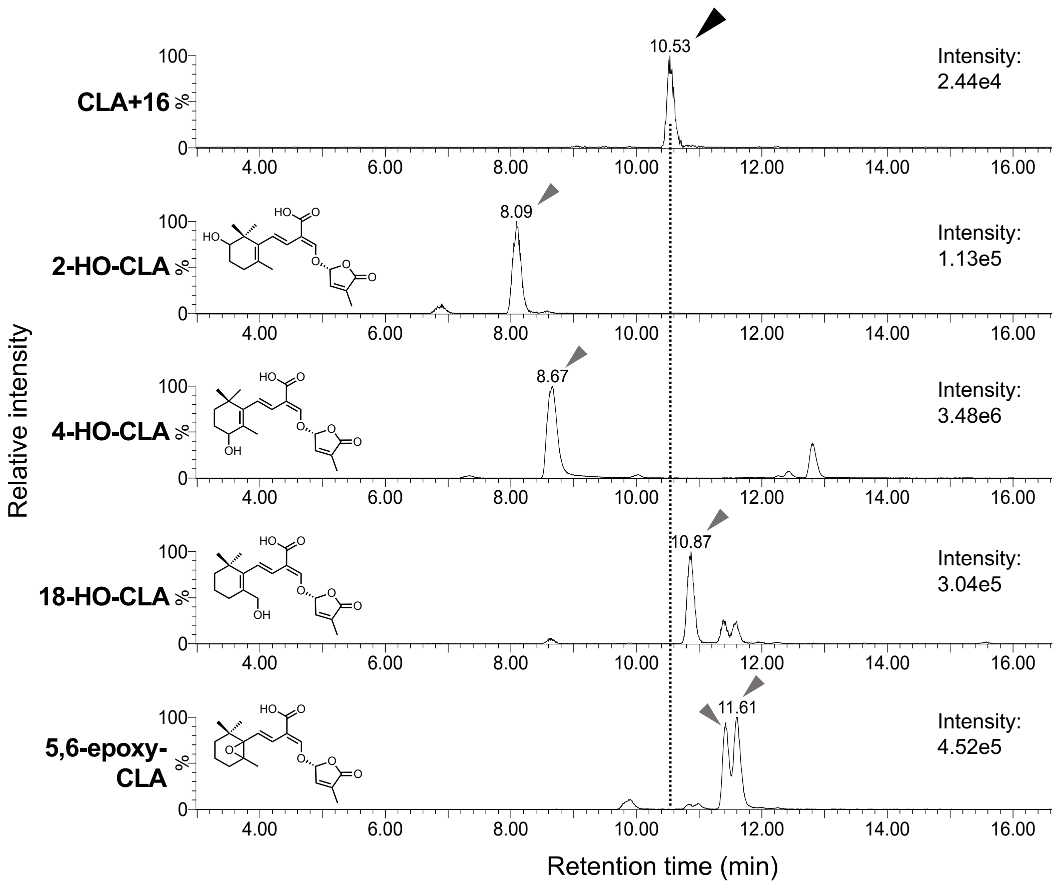


**Supplementary Fig. S5.** Comparison of retention times of the CLA+16 compound with other CLA derivatives in LC-MS/MS analysis.

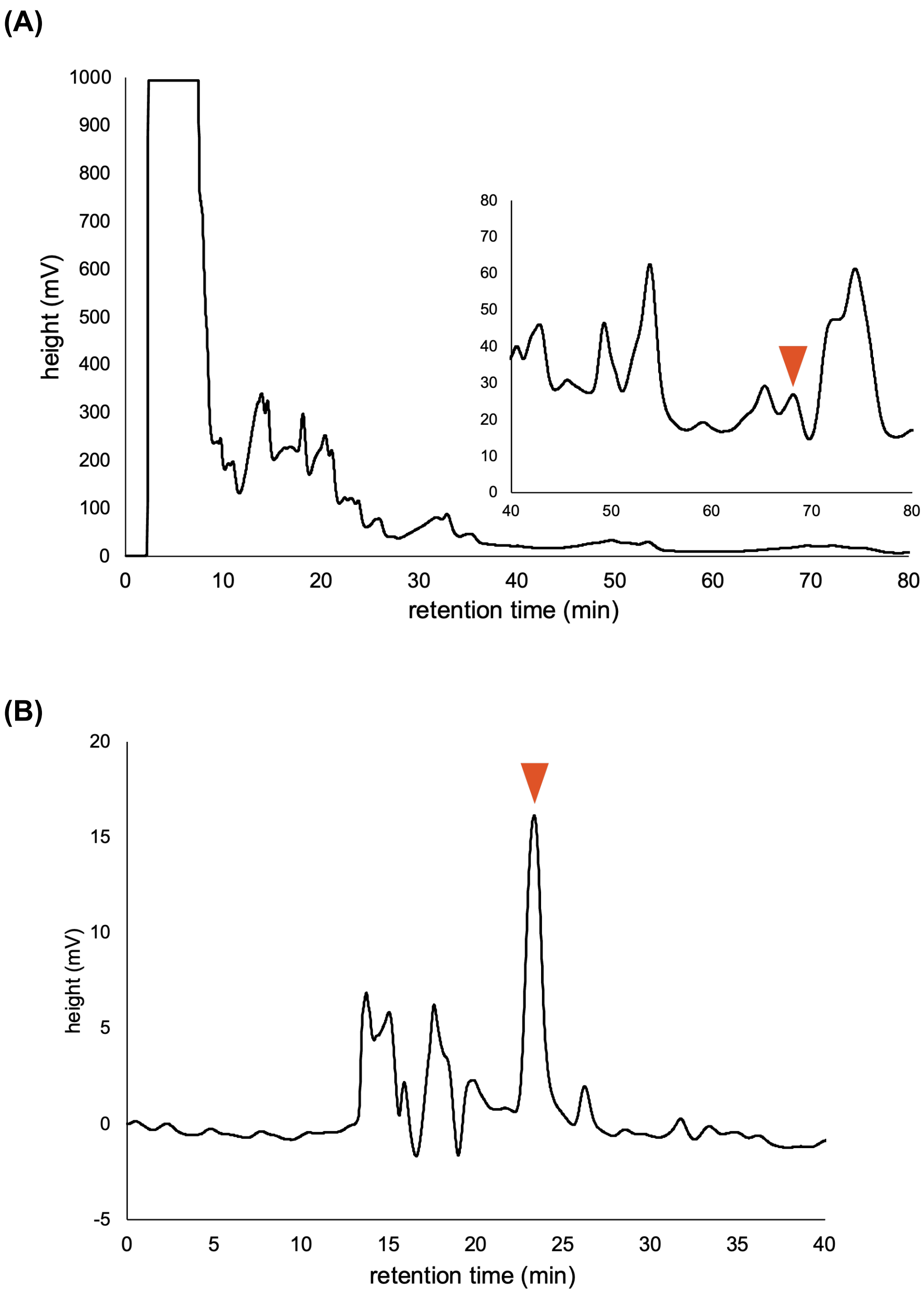


**Supplementary Fig. S6.** Chromatographic purification of the CLA+16 compound. The CLA+16 compound was first purified using an ODS column (A) and subsequently fractionated using a chiral column (B). Arrows indicate the target compound, CLA+16.

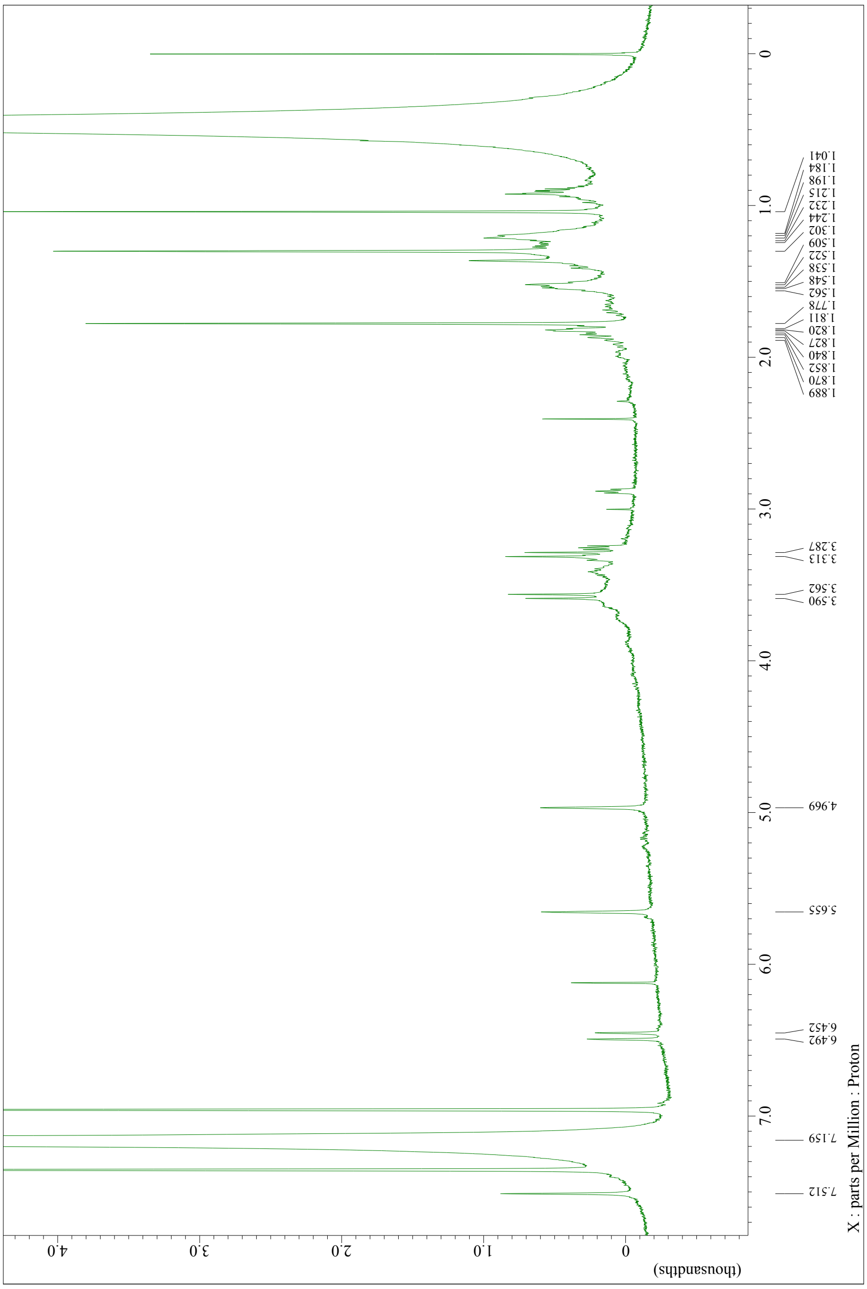


**Supplementary Fig. S7.** $^{1}$H-NMR spectrum of the CLA+16 compound identified as 16-HO-CLA.

**Supplementary Table S1** ¹H-NMR spectroscopic date of CLA+16 and CLA.

| No. | CLA+16 produced by CYP722A1 ($C_6D_6$) $\delta^1$H (mult., *J* Hz) | CLA ($C_6D_6$) $\delta^1$H (mult., *J* Hz) |
|---|---|---|
| 2 | 1.19 (*m*) | 1.28 (*m*) |
| 3 | 1.53 (*m*) | 1.54 (*m*) |
| 4 | 1.85 (*m*) | 1.89 (*t*) |
| 5 | | |
| 6 | | |
| 7 | * | 7.28 (*d*, 16.0) |
| 8 | 6.43 (*d*, 16.4) | 6.53 (*d*, 16.5) |
| 9 | | |
| 10 | 7.51 (*s*) | 7.54 (*s*) |
| 11 | 4.97 (*s*) | 4.96 (*s*) |
| 12 | 5.66 (*s*) | 5.61 (*s*) |
| 13 | | |
| 14 | | |
| 15 | 1.78 (*s*) | 1.81 (*s*) |
| 16a | 3.29 (*d*, 11.0) | 1.15(*s*) |
| 16b | 3.58 (*d*, 11.0) | |
| 17 | 1.04 (*s*) | 1.13 (*s*) |
| 18 | 1.30 (*s*) | 1.36 (*s*) |

* Overwrapped with the solvent signals.
